# Crippling the bacterial cell wall molecular machinery

**DOI:** 10.1101/607697

**Authors:** Allison H. Williams, Richard Wheeler, Ala-Eddine Deghmane, Ignacio Santecchia, Francis Impens, Paulo André Dias Bastos, Samia Hicham, Maryse Moya Nilges, Christian Malosse, Julia Chamot-Rooke, Ahmed Haouz, William P. Robins, Muhamed-Kheir Taha, Ivo Gomperts Boneca

## Abstract

Lytic transglycosylases (LT) are redundant enzymes that play a critical role in peptidoglycan (PG) recycling and metabolism. LT(s) role in cell wall-modifying complexes and usefulness as antimicrobial drug targets remain elusive. We determined at high-resolution a structure of the membrane-bound homolog of the soluble LT from *Neisseria* species with a disordered active site helix (alpha helix 30). Alpha helix 30 is crucial for binding PG during catalysis^1^. Here we show using an alpha helix 30 deletion strain that LT (LtgA) determines the integrity of the cell wall, participates in cell division and separation, and can be manipulated to impair the fitness of the human pathogen *Neisseria meningitidis* during infection. Characterization of *ltgA* helix deleted strain interactome identified glycan chain remodeling enzymes whose function appear to be modulated by LTs. Targeting LTs can disrupt the PG machinery, which is fatal for the bacterium, a new approach for antibiotic development.

## Introduction

Lytic transglycosylases (LTs) degrade peptidoglycan (PG) to produce *N*-acetylglucosamine (GlcNAc)-1,6-anhydro-*N*-acetylmuramic acid (MurNAc)-peptide (G-anhM-peptide), a key cytotoxic elicitor of harmful innate immune responses^2^. LTs have been classified into four distinct families based on sequence similarities and consensus sequences. LTs belonging to family 1 of the glycoside hydrolase (GH) family 23 share sequence similarity with the goose-type lysozyme^3^. Family 1 can be further subdivided into 5 subfamilies, 1A through E, which are all structurally distinct^3^. Despite the overall structural differences among LTs, their active sites, enzymatic activities and substrate specificities are fairly well conserved.

The crystal structure of the outer membrane lipoprotein LtgA, a homolog of Slt70 that belongs to family 1A of GH family 23 from the pathogenic *Neisseria* species, was previously determined at a resolution of 1.4 Å (Fig. 1a). ^4,5^. Briefly, LtgA is a highly alpha-superhelical structure consisting of 37 alpha helices (Fig. 1a). Although LTs have very diverse overall secondary structures, they exhibit similar substrate specificity and a preference for PG^6^. LtgA shares an overall weak sequence similarity with Slt70 (25%). However, the structural and sequence alignments of the catalytic domains of Slt70 and LtgA revealed absolute active site conservation^5^. The active site of LtgA is formed by ten alpha helices (α 28, 29, 30, 31, 32, 33, 34, 35, 36, 37), with a six-alpha-helix bundle (α 29, 30, 31, 32, 33, 34) constituting the core of the active site that firmly secures the glycan chain (Fig. 1a).

**Figure 1.**
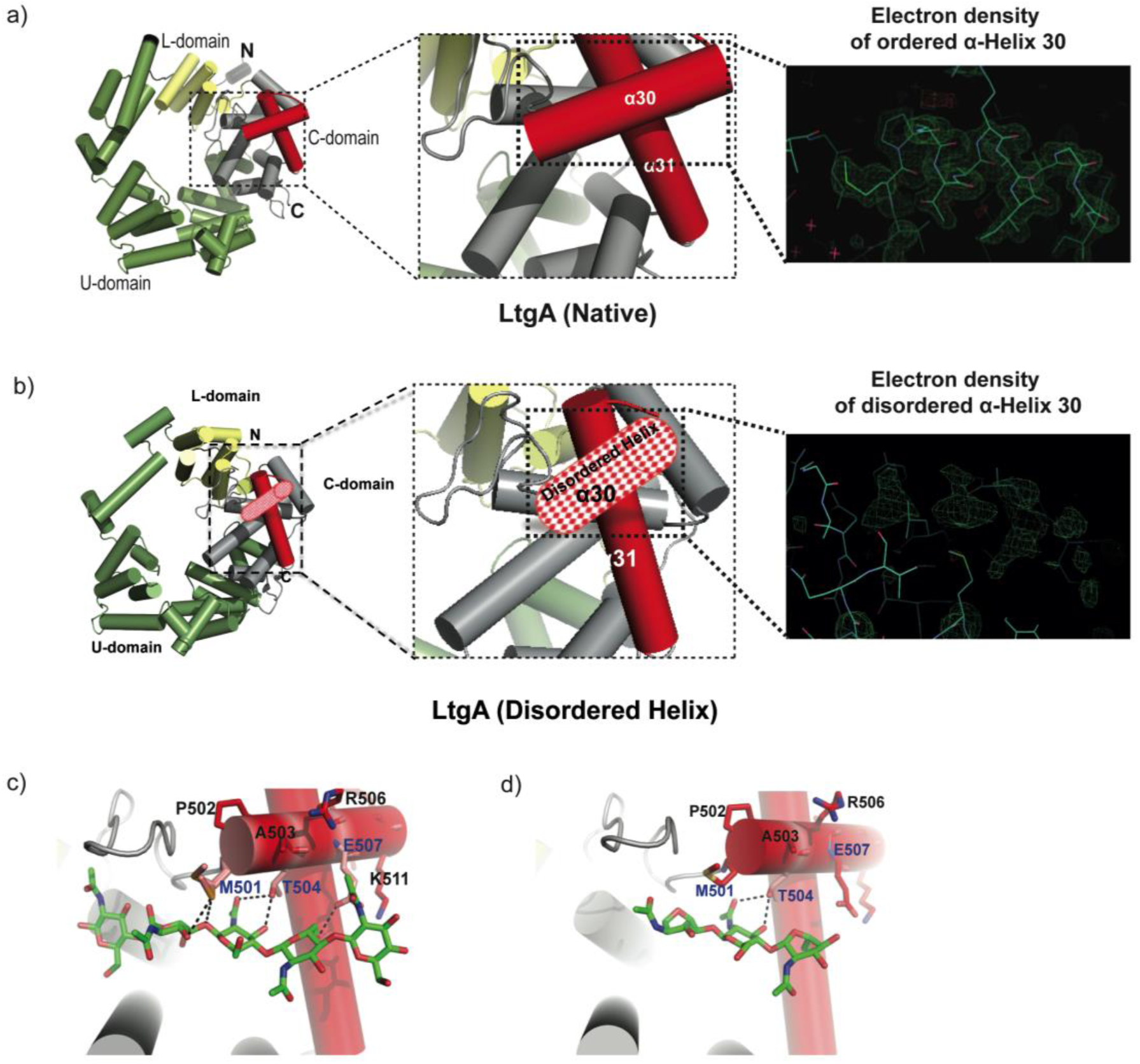
Molecular architecture of LtgA alpha helix 30 and contacts made with reaction intermediates. a) Native structure of LtgA. Ribbon model of LtgA displaying a helical structure consisting of 37 alpha helices. LtgA consists of three domains: A C-domain (gray and red), which houses the putative catalytic domain, and the L (yellow) and U (green) domains, which are of unknown function. A long N-terminal extension interacts with the L-domain, which closes the structure (PDB ID: 5O29). Clear and consistent density for helix 30 was depicted by the Fo-Fc omit map (green) b) LtgA with a disordered conformation of helix 30. Clear and consistent density for helix 30 was absent as depicted by the Fo-Fc omit map (green) of helix 30 (PDB ID: 6H5F). c) LtgA plus trapped intermediates (chitotetraose and a GlcNAc sugar) (PDB ID: 5O2N). d) LtgA plus anhydro product (1,6-anhydro-chitotriose) (PDB ID: 5OIJ).

LTs utilize a single catalytic residue, either a glutamate or aspartate, which plays the role of an acid and then that of a base^7–11^. In our recent study, active LtgA was monitored for the first time in the crystalline state, and the residues involved in the substrate and product formation steps were identified. Globally, conformational changes occurred in three domains, the U, C and L domains, between native LtgA and LtgA bound to the product^5^. Substantial conformational changes were observed in the active site, for example, during the product formation step, the active site adopted a more open conformation^5^.

Many Gram-negative bacteria have multiple and redundant LTs; for example, *Escherichia coli* has eight (MltA, MltB, MltC, MltD, MltE, MltF, MltG and Slt70), and *Neisseria* species encodes 5 (LtgA, LtgB, LtgC, LtgD, and LtgE), suggesting that LT activity is vital to the life cycle of the cell^12^. Because the activity of LTs is redundant, the loss of one or more LTs in *E. coli* leads to no observable growth defects. When genes for 6 LTs were deleted from *E. coli*, a mild chaining phenotype was observed^13^. However, despite lack of strong observable phenotypic changes, it has been suggested that LTs may have well-defined roles in the cell. For example, the deletion of *ltgA* and *ltgD* in *Neisseria gonorrhoeae* eliminates the release of cytotoxic PG monomers suggesting the activity of LtgA and LtgD are redundant. Moreover, LtgA primarily localizes at the septum, indicating a role in the divisome machinery; whereas, LtgD is distributed along the entire cell surface^14^.

The activities of LTs are known to be inhibited by β-hexosaminidase inhibitors (for example, NAG-thiazoline); bulgecins A, B and C; and PG-*O*-acetylation^4,10,15,16^. PG-*O*-acetylation^17^ is a process that allows pathogenic bacteria to subvert the host innate immune response^18,19^. It should be noted that many Gram-positive and Gram-negative bacteria *O*-acetylate their PG, with a few notable exceptions such as *E. coli and Pseudomonas aeruginosa*^20^. Peptidoglycan *O*-acetylation prevents the normal metabolism and maturation of PG by LTs^21^. Ape1, a PG de *O*-acetylase, is present in *Neisseria* species and generally in Gram-negative bacteria that *O*-acetylate their PG. Ape1 catalyzes the hydrolysis of the *O*-acetyl modification specifically at the sixth carbon position of the muramoyl residue, thus assuring the normal metabolism of PG by LTs^17,22,23.^

LTs are an integral part of the enzymatic PG machinery and forms protein complexes with other members of the PG biosynthetic apparatus, such as PBPs^6,12,24–27^. Most notable are the interactions between Slt70 and PBPs 1b, 1c, 2 and 3^28^. PBPs are essential for bacterial cell wall synthesis and are required for proliferation, cell division and the maintenance of the bacterial cell structure. Previously, PBPs were thought to be primarily responsible for the polymerization of PG. Recently, RodA, a key member of the elongasome, and a shape, elongation, division and sporulation (SEDS) protein family member was shown to be a PG polymerase. RodA functions together with PBP2 to replicate the transglycosylase and transpeptidase activities found in bifunctional PBPs^29–31^. SEDS proteins are widely distributed in bacteria and are important in both the cell elongation and division machinery. *Neisseria* species such as *N. gonorrhoeae* and *N. meningitidis* are coccoid in shape and lack an elongation machinery. Therefore, these species incorporate new PG through complex interactions in the divisome. Both *N. gonorrhoeae* and *N. meningitidis* have five PBPs, namely, PBP1, PBP2, PBP3, PBP4 and PBP5. PBP1 and PBP2 are homologous to *E. coli* PBP1a and PBP3, while the *Neisseria* PBP3 and PBP4 are homologous to *E. coli* PBP4 and PBP7^32^. PBP5 in both *E. coli* and *Neisseria* species are both predicted carboxypeptidase^33^.

FtsW, a RodA homolog and a key component of the divisome machinery, forms a complex with FtsI (PBP3), which has been shown to interact with PBP1b, FtsN and other proteins of the divisome^34^. The FtsW-PBP3 complex shares similar interacting regions with the RodA-PBP2 complex, and is the confirmed PG polymerase of the divisome^35^.

Previous work by our group and others have demonstrated that PBPs and LTs can be targeted in a combined antibiotic regimen that could counter antibiotic resistance^36^, highlighting the possibility of simultaneously inhibiting LTs and their binding partners, such as PBPs, to achieve a synergistic antibiotic effect. Here, we reveal the near-atomic-resolution crystal structure of a native version of LtgA with a disordered active site alpha helix. The deletion of LtgA alpha helix 30 affected bacterial growth, disrupted cell division and daughter cell separation, compromised the integrity of the cell wall and PG composition and diminished bacterial fitness or virulence in a mouse infection model. We established the first PG-centric LT interactome study that identified protein partners directly associated with PG assembly (the primary focus of this study) that could be responsible for the observed pleitropic phenotypes. A small hub of glycan strand PG-associated enzymes, that are possibly controlled, functionally regulated, and stabilized by LTs was identified. This study demonstrates that individual LTs, despite their redundancy maybe essential and will likely guide future efforts on how to target LTs for future antibiotic development.

## Results

### Structure of LtgA with a disordered alpha helix 30

In the course of monitoring the LtgA reaction in the crystalline state, we captured a native version of LtgA with a distinctly disordered alpha helix 30 (Fig. 1b, movie 1). This represents a newly identified conformational state of LtgA whereby alpha helix 30 transitions from an ordered to a disordered state (Fig 1a-b). Interestingly, this disorder of alpha helix 30 did not affect the overall structural integrity of the active site (Fig. 1b, movie 1) because all the other helices making up the catalytic domain remained intact. Moreover, LtgA was already shown to be active in the crystalline state in our previous studies, although the molecular details of alpha helix 30 interactions with the ligand was not addressed (Fig. 1a-b, movie 1)^5^.

Alpha helix 30, with the sequence ^501^(MPATAREIAGKIGMD)^516^ (Fig. 1a-b, colored in light red), is structurally conserved among the closest homologs of LtgA, mainly, Slt’s, and other LTs such as MltE and MltC ^8,37–40^ (Supplementary Fig. 1). Alpha helix 30 clamps the glycan strand during catalysis (Fig. 1c, movie 1) and undergoes conformational changes to a more open conformation after product formation (Fig. 1d). Met 501 and Glu 507 of alpha helix 30 lose hydrogen-bonding contact with the ligand after product formation (Fig. 1d, movie 1). Consistent with the structural data showing the role of alpha helix 30 in substrate/product binding, a heterologously expressed and purified LtgA^Δ30^ showed severely diminished PG binding capabilities when compared to wild-type LtgA or mutants of residues involved in the catalytic mechanism or substrate binding (E481A, E580) of LtgA (Supplementary Fig. 2). This further emphasizes the potential critical structural role of alpha helix 30 in the function of LtgA and consequently in the proper metabolism of the PG.

### The functional role of alpha helix 30

Given the important structural role of LtgA alpha helix 30, we investigated its functional role *in vivo* by engineering the following constructs in *N. meningitidis*: i) an LtgA knockout strain (Δ*ltgA*), ii) a knockout strain complemented at an ectopic locus on the meningococcal chromosome with the wild-type gene (Δ*ltgA*^*ltgA*^), or iii) complemented with alpha helix 30 deletion (Δ*ltgA*^*ltgA*Δ30^). Similar to other LTs, a complete deletion of the *ltgA* gene from the chromosome did not affect the growth of the bacteria (Fig. 2a)^41^. Interestingly, the strain with *ltgA* lacking the alpha helix 30 coding sequence exhibited severely reduced growth (Fig. 2a), with an exponential phase growth rate (0.059 h^−1^±0.012) significantly different from that of the wild-type or Δ*ltgA*^*ltgA*^ strain (0.72 h^−1^±0.15 or 0.21 h^−1^ *±*0.043, respectively) based on the calculated slopes of the growth curves (*p*<0.0001) (Fig. 2a).

**Figure 2.**
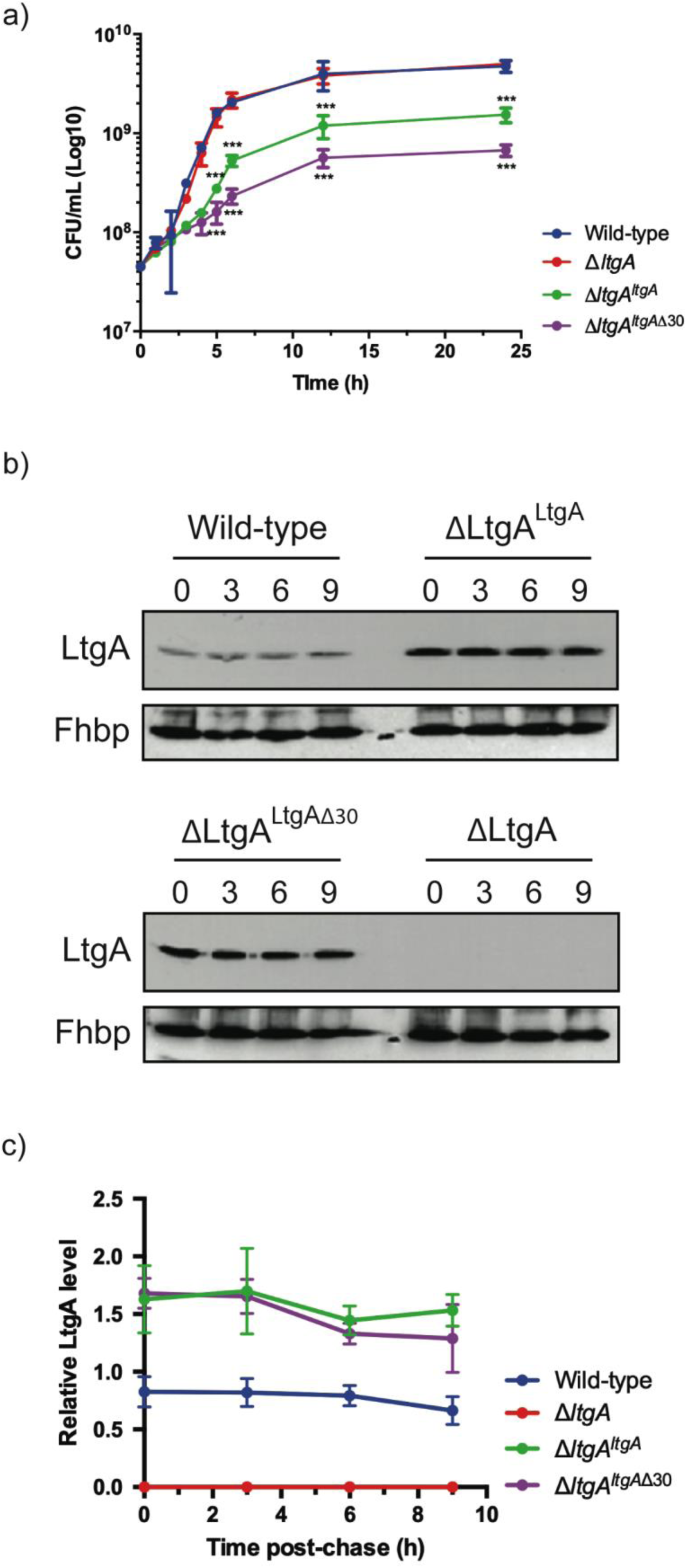
The LtgA helix 30 mutant leads to a growth deffect and is stable. a) Growth kinetics of *N. meningitidis* wild-type, *ΔltgA, ΔltgA*^*ltgA*^ and *ΔltgA*^*ltgA*Δ30^. Data represent three independent experiments with *n* = 3. b) Exponentially grown bacteria were treated with chloramphenicol (2 μg/ml) to block protein synthesis and survey the stability of LtgA for the indicated periods of time (in hours). Immunoblot was performed probing with anti-LtgA antibody. The expression of the outer membrane protein Fhbp was used as an loading control. c). The levels of LtgA over the time were analysed and ploted as a stability curve by quantifying the band intensities using ImageJ software. For each strain, the LtgA intensity at time zero is referred to as 100%, while the simultaneously Fhbp was used for loading control.

To exclude concerns about LtgA stability and to confirm that LtgA^Δ30^ continued to be expressed, the degradation of LtgA across all four strains was examined by western blotting following *de novo* protein synthesis in the lysates of bacteria harvested at various time points after incubation with chloramphenicol (Fig. 2b-c). The Δ*ltgA* knockout mutant strain did not exhibit any signal (Fig. 2b-c). The level of LtgA in the complemented strains was slightly higher than that in the wild-type strain at t_0_, possibly because the transcription of *ltgA* was controlled by a stronger promoter than that in the parental strain. After the addition of chloramphenicol, LtgA appeared to be maintained at comparable levels in the wild-type and complemented strains, and the levels decreased slowly during the sampling period, as revealed by quantitative measurement of relative protein abundance using densitometry (*t*_1/2_ > 9 h) (Fig. 2b-c).

The promoter for *ltgA* has not yet been identified; therefore, *ltgA* was introduced in the chromosome of meningococcus and expressed under the control of a non-native promoter. Since Δ*ltgA*^*ltgA*Δ30^ exhibited reduced growth and this could be attributed to bacterial lysis or defects in cell division or cell separation, we examined all four strains using fluorescent microscopy (labelling the cell wall and intracellular DNA), and scanning electron microscopy (SEM). Despite the reduced growth of strain Δ*ltgA*^*ltgA*Δ30^, there was no physical evidence suggesting bacterial lysis. However, intriguingly in the Δ*ltgA*^*ltgA*Δ30^ strain, we observed, strong defects in cell separation and cell division, and the appearance of membrane stained extracellular material that were notably absent in the other three strains (Fig. 3 (*right panel*), Supplementary Fig. 3a-b). Additionally, SEM revealed large blebs on the surface of some of unseparated/undivided bacteria in the Δ*ltgA*^*ltgA*Δ30^ strain that were not observed in the other strains. A rather striking phenomenon is that the bacteria with blebs all had smooth surfaces that deviated from the normal rough surface appearance of *N. meningitidis* in the other strains (Fig 3 *(left panel)*, Supplementary Fig. 3a-b). We identified ghost cells, however this phenomenon was not as pervasive as other abnormalities (Fig 3 *(left panel)*).

**Figure 3.**
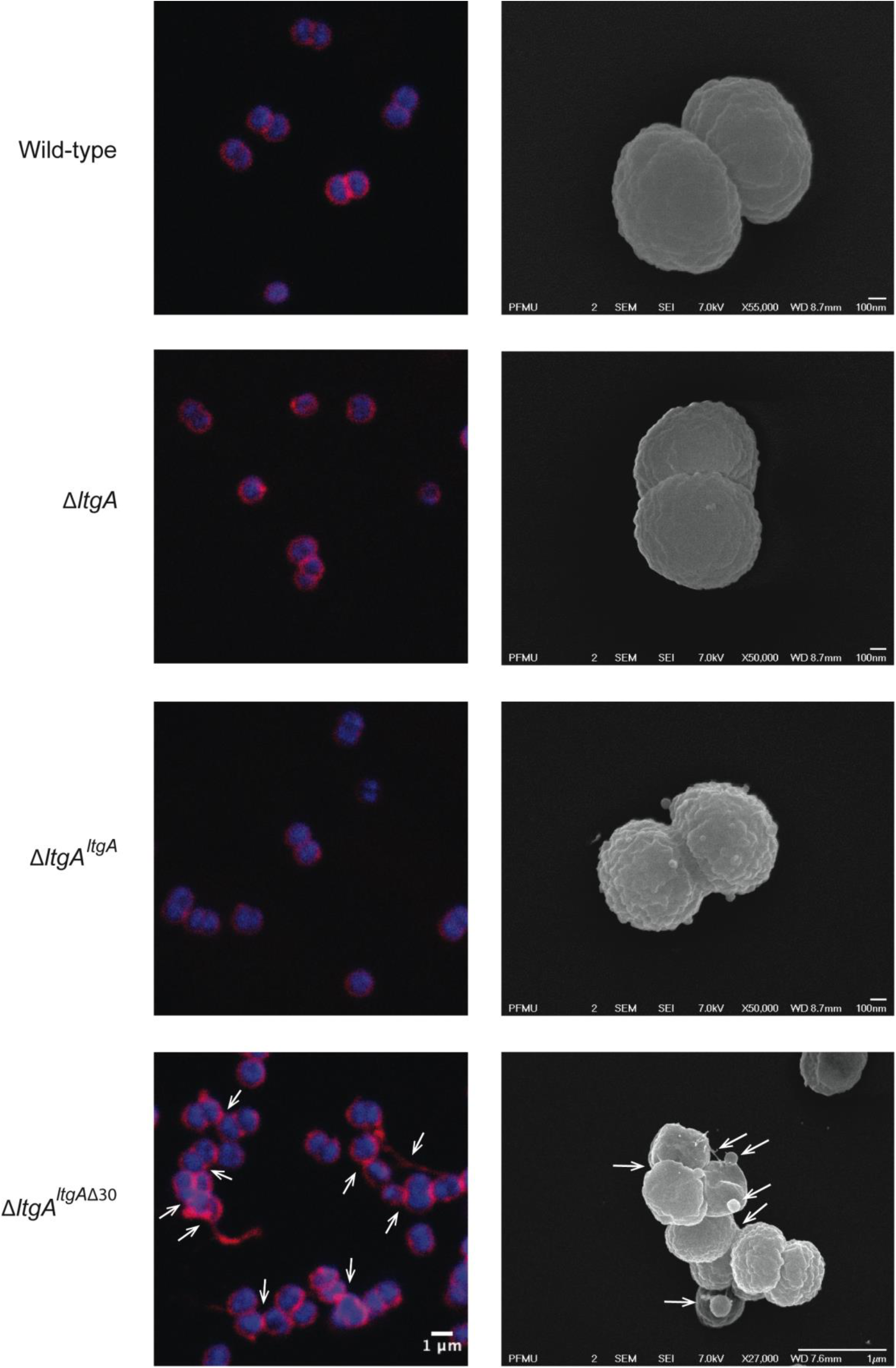
The LtgA helix 30 mutant shows morphological abnormalities. Morphological differences between strains of wild type and mutant lytic transglycosylases were determined by fluorescent microscopy (*right panel*) and scanning electron microscopy (SEM) (*left panel*). White arrows in the images of *ΔltgA*^*ltgA*Δ30^ strain (*right panel*) points to cells defective in division and separation, as well as extracellular material. White arrows in the left panel points to, irregular cell surfaces, high molecular weight blebs (not observed in other strains), asymmetrical diplococci, and ghost cells. (See Supplementary Fig. 3a-b for other images detailing additional morphological abnormalities)

### LtgA is responsible for maintaining the structural integrity of PG

We examined the PG profiles of wild-type *N. meningitidis*, Δ*ltgA*, Δ*ltgA*^*ltgA*^ and Δ*ltgA*^*ltgA*Δ30^, to explore whether the integrity of the PG composition of Δ*ltgA*^*ltgA*Δ30^ strain was maintained. No notable differences were observed among the wild-type, Δ*ltgA* and Δ*ltgA*^*ltgA*^ strains (Fig. 4, Supplementary Fig. 4, Supplementary table 1). However, the PG of the Δ*ltgA*^*ltgA*Δ30^ strain was found to be markedly hyperacetylated when compared to that of the other strains, with a 102% increase in the amount of acetylated GlcNAc-anhMurNAc-tetrapeptide (GM*4), a 39% increase in acetylated GlcNAc-anhMurNAc-tetrapeptide crosslinked with GlcNAc-MurNAc-tetrapeptide (GM*4-GM4), and a 46% increase in doubly acetylated di-GlcNAc-anhMurNAc-tetrapeptide (GM*4-GM*4) (Fig. 4, Table 1). A 22% increase in the amount of GlcNAc-MurNAc tetrapeptide (GM4) was also observed, while the amounts of GlcNAc-MurNAc tripeptide (GM3) and GlcNAc-MurNAc pentapeptide (GM5) decreased by 33% (Fig. 4, Table 1). Overall, there was a marked increase in the amounts of acetylated PG monomers and dimers.

**Figure 4.**
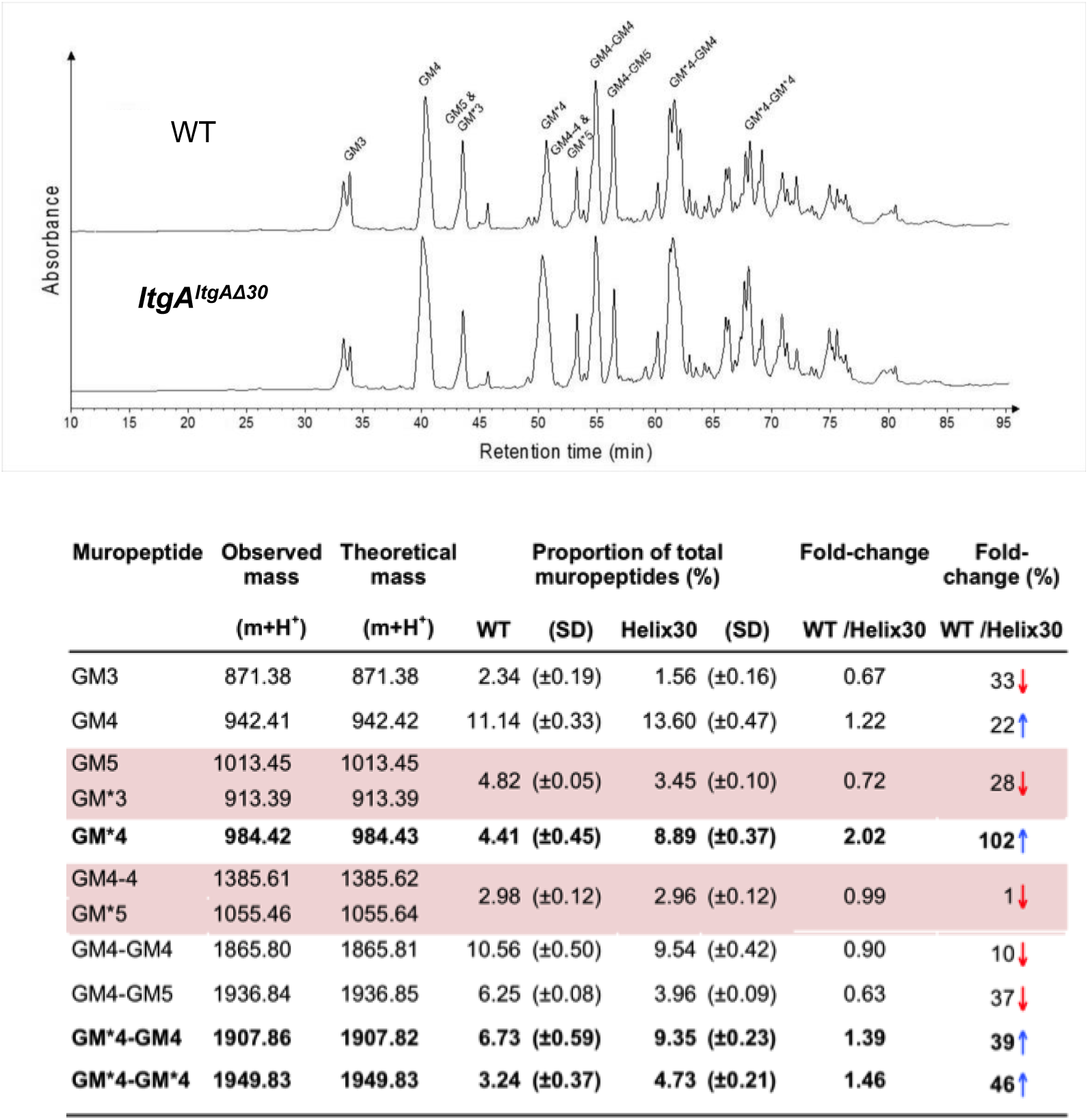
Muropeptide composition of PG isolated from wild-type and Δ*ltgA*^*ltgAΔ30*^. Purified PG was digested by muramidase mutanolysin, and the resulting muropeptides were reduced and then analyzed by LC/MS. The results were reproducible over 4 biological replicates. Peak identifications correspond to Table 1. Table 1. Muropeptides identified by mass spectrometry.* indicates O-acetylated MurNAc. Acetylated GM*4 is highlighted in bold. Multiple muropeptides coeluted as a single peak are shaded in pink. Red arrows indicate a decrease and blue arrows an increase in muropeptide abundance. The table displays the observed and theoretical masses and the proportion of total muropeptides.

### The protein-protein interactome of LtgA reveals binding partners involved in the PG machinery

To date, there have been no PG-centered omic studies on enzymes involved in cell wall biosynthesis. Deletion of the LtgA alpha helix 30 had a multifactoral impact on *Neisseria*, including impaired growth, defects in cell division or cell separation, and altered PG composition. Therefore, we decided to explore whether impaired LtgA activity could perturb the protein molecular environment of PG assembly.

To identify partners of LtgA potentially involved in PG and cell wall metabolism we utilized coimmunoprecipitation followed by high-performance affinity enrichment mass spectrometry using *ΔltgA*, Δ*ltgA*^*ltgA*^ and Δ*ltgA*^*ltgA*Δ30^ strains (Fig. 5a-b). These coimmunoprecipitation experiments were performed with each strain in triplicate (Fig. 5a-b). To our knowledge, this experiment is the first large-scale assessment of the protein-protein interactions of LTs. We then created a protein-protein interactome (PPI) network and, carried out a Clusters of Orthologous Groups (COG) analysis to classify the proteins identified in the PPI network for the Δ*ltgA*^*ltgA*Δ30^ strain (Supplementary Fig. 5). We discovered that the proteins identified in this network belong to 2 main functional categories: i) cell wall/membrane/envelope biogenesis and ii) intracellular trafficking, secretion, and vesicular transport (Supplementary Fig. 5). Intriguingly, the interactors of LtgA^Δ30^ identified in the Δ*ltgA*^*ltgA*Δ30^ mutant strain showed enrichment of peptides from known members of the PG synthesis machinery, that along with LtgA utilize the PG glycan strand as substrate (PBP1a, and the *E. coli* MltD homolog LtgE and LtgD a homolog of *E. coli* MltB) (Fig. 5,a-b, Supplementary table 2).

**Figure 5.**
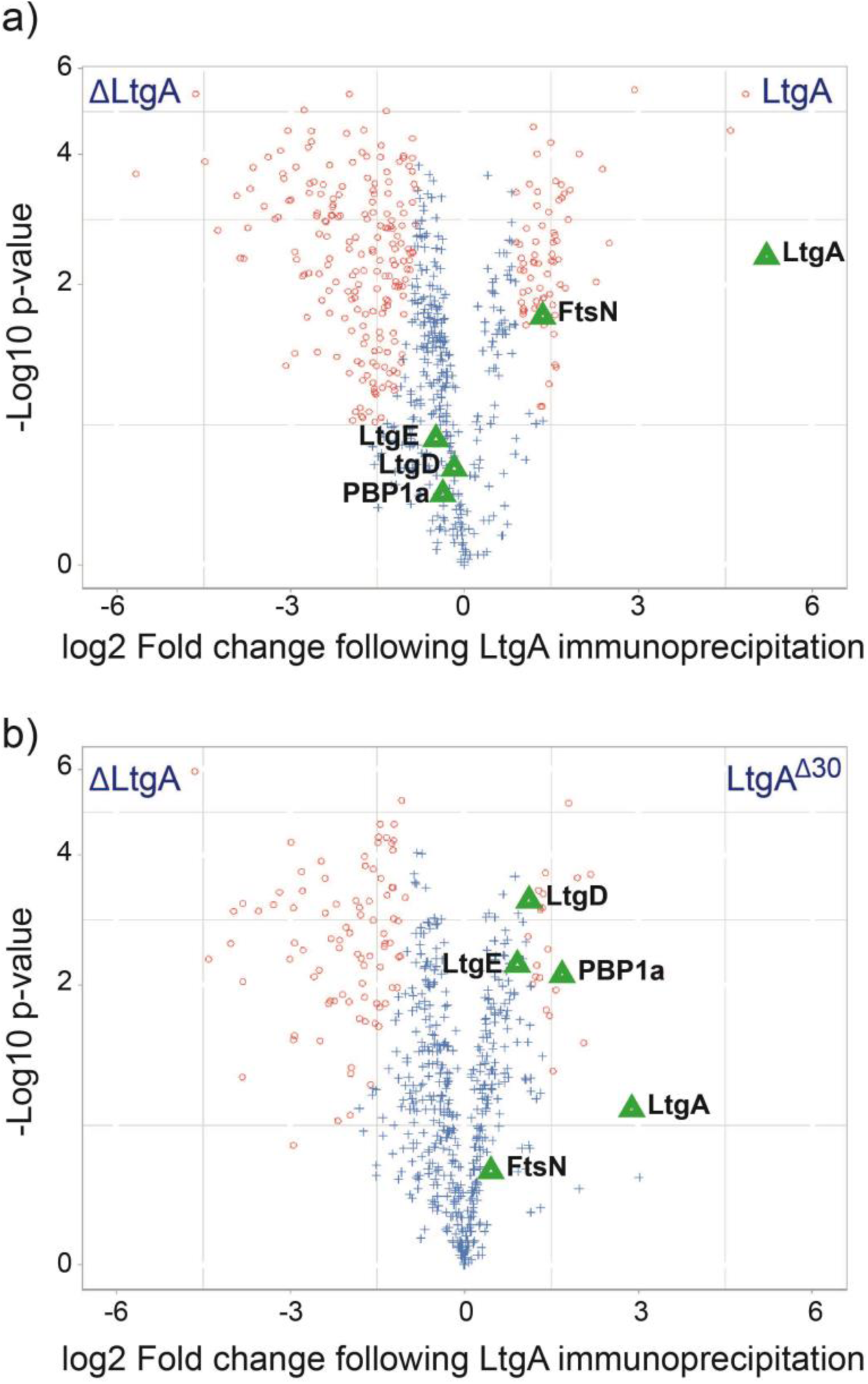
Protein-protein interactome (PPI) centered on LtgA. a) Volcano plots of proteins enriched in Δ*ltgA*^*ltgA*^ versus Δl*tgA* strains. b) Volcano plots of proteins enriched in Δ*ltgA*^*ltgA*Δ30^ versus Δl*tgA* strains. Coimmunoprecipitation experiments were performed with each strain in triplicate. The strains utilized were Δl*tgA*, Δ*ltgA*^*ltgA*^ and Δ*ltgA*^*ltgA*Δ30^. LC-MS/MS was used to identify proteins. Volcano plots depict significance versus fold change on the y-and x-axes, respectively. Data points with low (significant) p-values appear closer to the top of the plot (for p-values see Supplementary table 2). Changes in both directions (enrichment and lack of enrichment) appear equidistant from the center on the x-axis.

As expected, LtgA was found to be significantly enriched in the Δ*ltgA*^*ltgA*^ strain compared with the *ΔltgA* strain. No other enzymes directly involved in PG assembly were enriched in the Δ*ltgA*^*ltgA*^ strain, however, FtsN an essential divisome protein, previously shown to activate septal PG synthesis and constriction of the cell was enriched (Fig. 5a-b, Supplementary table 2). In contrast, the deletion of a single alpha helix led to an enrichment of enzymes known to be involved in PG synthesis (PBP1a, LtgE and LtgD) (Fig. 5a-b, Supplementary table 2).

### Validation of LtgA interactors

Since increased levels of *O*-acetylated and non-crosslinked PG subunits were the most striking phenotypes of the Δ*ltgA*^*ltgA*Δ30^ mutant strain, we used gel filtration binding studies to identify and confirm potential interactors of LtgA that could be responsible for the observed aberrations in the composition of the PG. Ape1 the PG-de-*O*-acetylase (not significantly enriched in the co-immunoprecipitation studies) and other identified glycan strand synthesis or modifying enzymes (LtgE, and PBP1a) was chosen for further validation (Fig. 6a-e). PBP2 (FtsI) of *Neisseria* species is a homolog of PBP3 of *E. coli*, and in complex with FtsW is the putative PG polymerase of the divisome. PBP2 has transpeptidase activity and was not enriched in our screen, but was included in this study because PBP3 from *E. coli* was reported to form protein complex with Slt70 an homolog of LtgA^28^. We first purified Ape1, LtgA, LtgE, PBP1a and PBP2 following their heterologous expression in *E. coli* (Fig. 6a,c-e, Supplementary Figure. 7). Each enzyme was purified individually and then combined prior to their application to size-exclusion columns (Fig. 6a-e). LtgA formed a 100-kDa complex with Ape1 and a 150-kDa complex with PBP1a (Fig. 6a, c-e).

**Figure 6.**
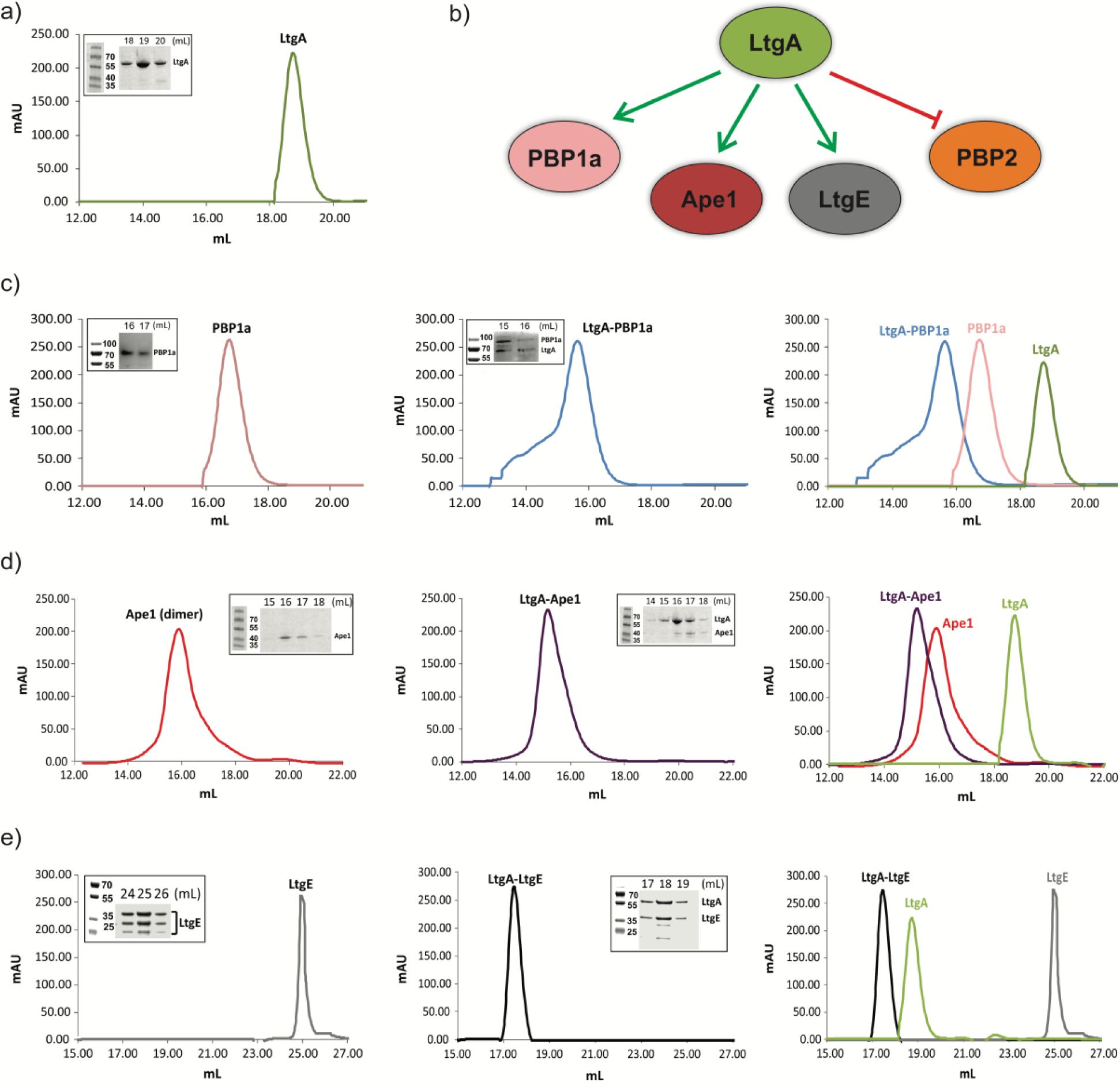
Validation of the LtgA PG-centric interactome. a) Chromatogram showing size exclusion analysis of LtgA (65 kDa, green). Each inserts represents an SDS-PAGE analysis of peak fractions containing proteins. Lanes are labeled with corresponding volumes. b) Summary of the validated LtgA PG-centric interactome. c, d, e) (*Left panel*) Chromatogram showing size exclusion analysis of LtgA interacting partners: PBP1a (85 kDa, pink), Ape1 (40 kDa, red) and LtgE N-terminal bacterial muramidase lysozyme-like domain (36 kDa, gray). (*Middle panel*) Chromatogram showing size exclusion analysis of the corresponding complexes with LtgA: PBP1a-LtgA (150 kDa, blue), Ape1-LtgA (105 kDa, purple) and LtgE-LtgA (101 kDa, black), respectively. Inserts are SDS-PAGE analysis of peak fractions containing LtgA protein complexes is shown in the inserts. (*Right panel*) Overlay of the chromatograms corresponding to purified LtgA, binding partners, and LtgA protein complexes.

LtgE is an approximately 72-kDa protein that is similar in molecular weight to LtgA (Fig. 6e). LtgE has two domains, namely, an N-terminal bacterial muramidase lysozyme-like domain and a LysM-like peptidoglycan-binding C-terminal domain (Fig. 6e). Because the full-length protein was recalcitrant to expression in *E. coli*, we successfully expressed instead the approximately 36-kDa N-terminal domain of LtgE which is prone to proteolytic degradation (Fig. 6e). However, the 36-kDa N-terminal domain of LtgE selectively binds LtgA to form an approximately 100-kDa complex with LtgA(Fig. 6e). Finally, the oligomeric PBP2 and monomeric LtgA was eluted as separate species, suggesting that there was no strong association between the two, validating the absence of significant enrichment in our coimmunoprecipitation study.

### The impact of protein complexes on peptidoglycan O-acetylation

Hyperacetylation of the PG was the most striking phenotype of the *ΔltgA*^*ltgA*Δ30^ strain, therefore we explored whether the normal function of Ape1 depends on LtgA or if Ape1 and LtgA work in concert enzymatically to de-*O*-acetylate the PG. To accomplish this, we examined the processivity of LtgA and Ape1 against acetylated PG from *N. meningitidis* or the activity of an Ape1 and Ape1 combined with LtgA toward 4-nitrophenyl acetate, a previously characterized substrate of Ape1 from *N. gonorrhoeae* that is not a substrate for LtgA ^42,43^.

In the presence of equimolar (1.2 μM) amounts of Ape1, LtgA degrades the PG more efficiently (Supplementary Fig. 7), this is consistent with previous studies that suggest *O*-acetylation blocks the function of LTs and lysozyme PG^17,22,23,44^. Surprisingly, in the absence of a common substrate and utilizing equimolar amounts (12 nM) of LtgA and Ape1, LtgA stabilizes and enhances the activity of Ape1. The reaction remained well within the linear range for 60 minutes when both enzymes were present, which was in stark contrast to Ape1, that showed less activity over the time course of 60 minutes. These data demonstrate that the enzymatic activities of LtgA and ApeI are enhanced reciprocally when functioning together in a complex (Supplementary Fig. 7a-b). It also appears that LtgA stabilizes and enhance the activity of Ape1.

### Role of alpha helix 30 in the virulence of N. meningitidis

In *N. meningitidis* and *N. gonorrhoeae*, the activity of LtgA and other LTs leads to a release of cytotoxic PG fragments, which are detected by the host and induce a Nod1-dependent inflammatory response^45–48^. Since the alpha helix 30-deleted strain of LtgA compromised the PG composition and caused enrichment of the protein partners involved in the PG machinery of enzymes, the functional role of the alpha helix was explored *in vivo* in a mouse infection model. For this purpose, we used transgenic mice expressing human transferrin as an experimental model that allows meningococcal growth by providing a human iron source during infection. The four *N. meningitidis* strains (wild-type, Δ*ltgA*^*ltgA*^, Δ*ltgA*^*ltgA*Δ30^ and Δ*ltgA*) were used to infect the mice by intraperitoneal injection. Two hours after infection, the four strains induced similar levels of bacteremia (Fig. 7a), suggesting that the strains were not defective in their ability to reach the bloodstream. The Δ*ltgA* strain appears to be cleared more slowly. However, the Δ*ltgA*^*ltgA*Δ30^ strain was cleared from the blood at a significantly faster rate than the other strains, exhibiting a 2-log difference in colony forming units (CFUs) at the 6-h time point compared to the wild-type strain (Fig. 7a). These results were also consistent with those at the cytokine production level, as the Δ*ltgA*^*ltgA*Δ30^ strain exhibited significantly decreased levels of IL-6 and KC (the functional murine homolog of human IL-8) 6 h after infection, while all the isolates exhibited similar levels 2 h after infection (Fig. 7b). Overall, the Δ*ltgA*^*ltgA*Δ30^ strain displays impaired fitness in the host, suggesting LtgA could play a key role in bacterial virulence.

**Figure 7.**
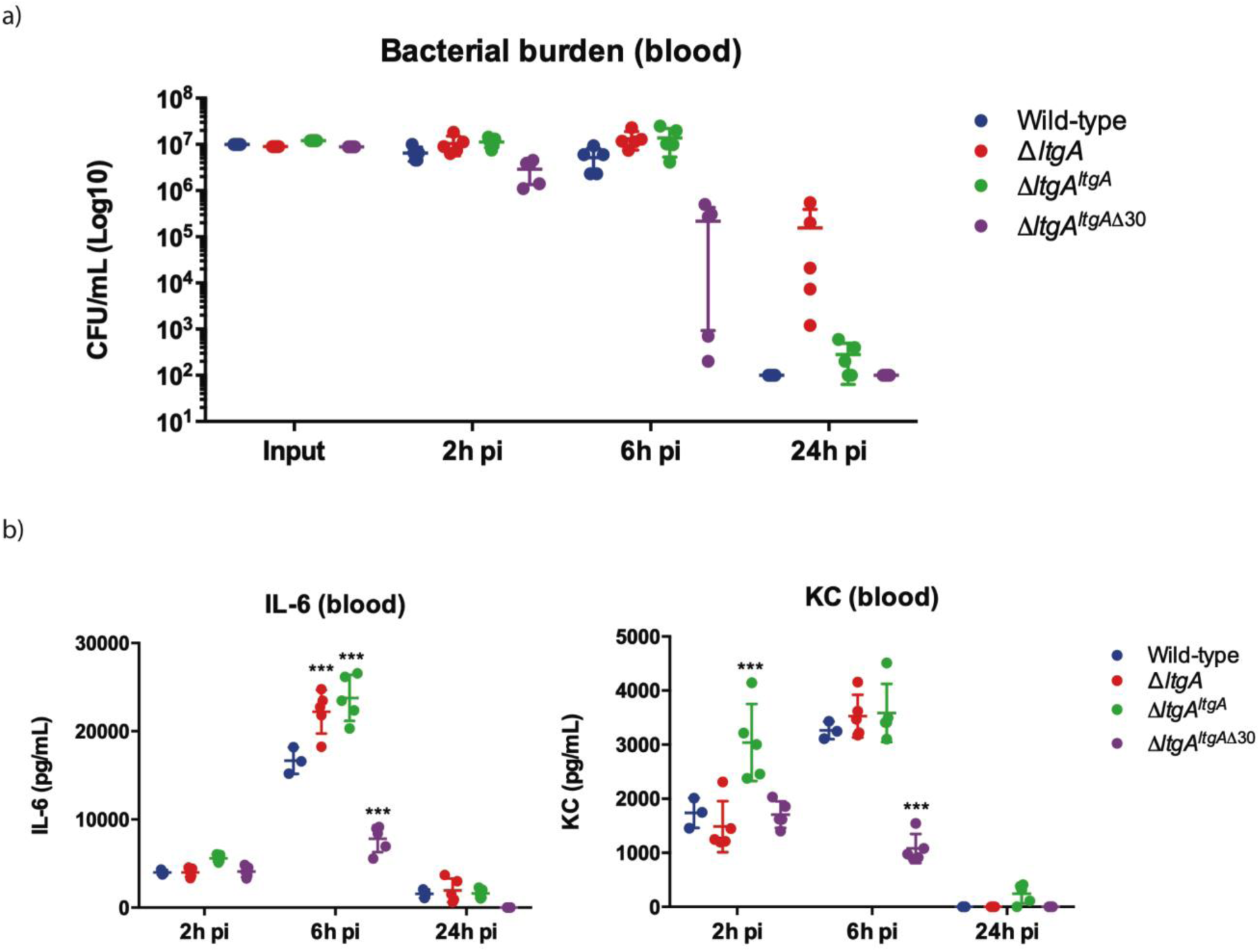
Helix 30 of LtgA plays a role in *Neisseria meningitidis* host adaptation and virulence. *N. meningitidis* wild-type, *ΔltgA, ΔltgA*^*ltgA*^ and *ΔltgA*^*ltgA*Δ30^ were administered to transgenic mice expressing human transferrin via intraperitoneal route. (a) Bacterial burden was determined by CFU enumeration in blood 2, 6 and 24 h pi. These data shows that the strain complemented with a deletion in helix 30 is cleared faster than the other strains. (b) Pro-inflammatory cytokine (IL-6) and chemokine (KC) profile in blood of infected mice was evaluated 2, 6 and 24 h post infection by ELISA. Δ*ltgA*^*ltgA*Δ30^ induced lower levels of inflammatory mediators production upon infection compared to the other strains. Data represents 3 independent experiments with *n* = 5.

## Discussion

Antibiotic resistance is recognized as an urgent global public health threat. One potential solution is to develop new treatment strategies that impact multiple cellular targets and consequently, circumvent the rise of antibiotic resistance. The bacterial cell wall is assembled by a number of enzymes, some of which are broadly categorized as PG polymerases, PG-modifying enzymes and PG hydrolases. The primary targets of β-lactams, clinically the most utilized antibiotics, are PBPs that are known to polymerize PG. The development of a combinatorial or single therapy that interferes with multiple cellular function of the PG machinery could define a new era in antibiotic development. This would be particularly relevant for *N. gonorrhoeae* infections, as no vaccines against this species are currently available, and highly resistant strains are on the rise. Indeed, a *N. gonorrhoeae* “superbug” has already been identified that does not respond to the usual treatment with β-lactams such as ceftriaxone^49^.

In this study, we identified a variant of LtgA with a disordered active site alpha helix 30, which is important for PG binding and for the catalytic mechanism of LtgA (Fig. 1b, movie 1). LTs are highly redundant enzymes, and when individual or multiple LTs are deleted, bacteria are known to proliferate normally because others LTs compensate for the loss of activity and/or function^45,50^. Interestingly, when 6 LTs were deleted from *E. coli*, only a mild chaining phenotype was observed^13^, however a mutant of *ltgC* with a 33-bp deletion at the 5′ end from *Neisseria* sp., showed defects in growth and daughter cell separation^51^. In our *ΔltgA*^*ltgA*Δ30^ strain, we observed significant defects in growth, cell division, cell separation, cell membrane irregularities, and fibrous and membranous extra cellular material, which were noticeably absent in the wild type, *ΔltgA*, and the Δ*ltgA*^*ltgA*^ strains (Fig. 2, 3, Supplementary Fig. 3a-b). LtgA has been shown to localize to the septum, but no phenotype associated with cell division or cell separation was previously reported. However, the expression of an LT with impaired function (LtgA^Δ30^) results in the perturbation of various processes that are essential for bacterial proliferation.

Having observed that the *ΔltgA*^*ltgA*Δ30^ strain, but not the *ΔltgA* strain, was defective in growth, cell separation, and cell division, we hypothesized that this pleiotropic phenotype could arise because the defective LtgA enzyme continues to participate in PG biosynthesis complexes (Fig. 3, Supplementary Fig. 3a-b). This could lead to aberrant activities by LtgA interaction partners, which compromise the fitness of the bacterium. We first analyzed the PG profiles of wild-type *N. meningitidis*, Δ*ltgA*, Δ*ltgA*^*ltgA*^ and Δ*ltgA*^*ltgA*Δ30^, and observed that Δ*ltgA*^*ltgA*Δ30^ strain was hyperacetylated and there was an increase in PG monomers (Fig.4, Table 1). To identify protein assemblies that could be responsible for these abnormalities, we examined the LtgA interactome using co-immunoprecipitation and gel filtration chromatography. We identified four enzymes, PBP1a, Ape1, LtgE (an *E. coli* MltD homolog) and LtgD (an *E. coli* MltB homolog) that interact with LtgA (Fig. 5a-b, 6a-e, and Supplementary table 2). Incidentally, Ape1, LtgE and LtgD similar to LtgA are PG glycan strand modifying enzymes and one of the activities of the bifunctional PBP1a is to polymerize the PG glycan strand. Interestingly, PBP2 with known transpeptidase activity did not interact with LtgA using gel filtration binding studies. Together (LtgA, LtgE, LtgD, Ape1, PBP1a,) appear to form the nucleus of the PG glycan strand remodeling machinery. Interestingly, consistent with our observation, PBP1a, LtgE, and LtgA all appear to have co-evolved because they are co-conserved in all the proteobacteria that were surveyed (Supplementary Fig. 8). It should be noted that the betaproteobacteria have two slightly diverged PBP1a genes, but this was not the case for *Neisseria* (Supplementary Fig. 8). On the other hand, Ape1 was exclusively found in *Neisseria, Kingella, Snodgrassell*a, *Morococcus, Azovibrio*, and one isolate of *Burkholderia ubortensis*, suggesting Ape1 was potentially acquired later by lateral gene transfer (Supplementary Fig. 8).

The hyperacetylation of the PG was the most striking phenotype. The interaction of LtgA with Ape1 is not completely unexpected because Ape1, a PG-de-*O*-acetylase, removes the *O*-acetyl group from the C6-hydroxyl position of the glycan strand of *O*-acetylated PG and ensures the proper metabolism of the PG by LTs (LtgA, LtgD or LtgE and others)^17,22,23,44^. We hypothesized that LtgA stablizes the function of Ape1 and potentially the hub of glycan chain targeting enzymes that are known to be crucial for the proper metabolism of the PG (Fig. 6a-e). To test this, we used 4-nitrophenyl acetate, a known substrate of Ape1 but not LtgA. Surprisingly, LtgA stabilizes the activity and function of Ape1 well past the normal range (1 hr) of most *in vitro* enzymatic reactions, giving clear evidence that LtgA could orchestrate the activity, stability and function of Ape1 (Supplementary Fig. 7). Our study revealed an intimate relationship between the LtgA and Ape1, ie. LtgA stabilizes or regulates the function of Ape1 and an impaired LtgA appears to poison the PG machinery with devastating effects toward the survival of *Neisseria*. Interestingly, the trapped complexes represent sub-complexes that are a part of the known enzymatic cell wall glycan strand machinery (Fig. 5a-b and Fig. 6a-e). These results provide additional insight into the coordination of the activities of PG-associated enzymes during cell wall biosynthesis.

A rather surprising discovery from the interactome study was that LtgA interacts with LtgE (Fig. 5a-b). To our knowledge, this study is the first to show that LTs form protein complexes with each other. LTs can be categorized as enzymes with endolytic or exolytic activities. Although the activities of LTs are known to be redundant, the fact that these enzymes can coordinate their activities in a complex was previously unknown. Both the endolytic and exolytic activities of LTs are needed for the recycling of PG or for the release of cytotoxic PG monomers. The endolytic enzyme (LtgE) cleaves middle of the glycan strands, and the exolytic enzymes (LtgA or LtgD) cleave the end of the PG strands^1^. Specifically, deletion of *ltgA* and *ltgD* genes resulted in a major decrease in the production of the cytotoxic PG monomers, while *ltgE* deletion did not affect PG monomer release^45,48^.

We also demonstrated that the bifunctional PBP1a interacts directly with LtgA (Fig. 6c). PBP1a is the first identified PG polymerase and the current major target of known antibiotics (Fig. 6c). In addition, we observed an enrichment of FtsN, a cell division protein that activates septal PG synthesis and constriction of the cell, in the strain expressing native LtgA^52–55^. Bacterial two-hybrid studies have shown that FtsN interacts with the *E. coli* divisome machinery, including the PG polymerase complex, FtsW-PBP3 (*Neisseria*, PBP2), FtsQ, and FtsA^34,55–57^. The nuclear magnetic resonance structure of FtsN of *E. coli* reveals that it is a bitopic membrane protein with a small cytoplasmic portion and a large domain on the periplasmic face^58^. Future studies could establish whether, the periplasmic portion of FtsN interacts either directly or indirectly with LtgA and other periplasmic proteins. LtgA is localized at the septum^15,24^, and FtsN activates septal PG synthesis at the cell division site. Therefore LtgA could be connected to the divisome machinery through FtsN. This could be one of the explaination as to why an impaired LtgA appears to disrupt the normal course of cell division and cell separation.

Finally, to understand what role a defective LtgA that interferes with the normal function of the PG machinery plays in the pathogenesis of *N. meningitidis* we used a mouse infection model and showed Δ*ltgA*^*ltgA*Δ30^ strain of type *N. meningitidis* was cleared from the blood at a significantly faster rate than the wild-type, Δ*ltgA*^*ltgA*^ or Δ*ltgA* strains. Consistent with the virulence phenotype, and in comparison to the other three strains, the Δ*ltgA*^*ltgA*Δ30^ exhibited significantly decreased levels of IL-6 and KC, 6 h post-infection (Fig. 7b) indicating a general loss of fitness of the helix-30 deleted mutant. Unexpectedly, the Δ*ltgA* strain cleared the blood stream at a slower rate than the wild-type, Δ*ltgA*^*ltgAΔ30*^ *or* the Δ*ltgA*^ltgA^. However, the levels of the cytokine IL-6 and KC were comparable to that of the wild-type or the complemented strain. A deletion of both *ltgA* and *ltgD* in *N. gonorrhoeae* eliminates the release of cytotoxic PG monomers^14,59^. In our study, we deleted only *ltgA* and therefore the expression of LtgD continues to release soluble monomeric muropeptides, that could drive inflammation, while Δ*ltgA*^*ltgA*Δ30^ has a larger defect in the PG machinery. Additionally, the Δ*ltgA*^*ltgA*Δ30^ strain is hyperacetylated, suggesting Ape1 the PG de *O-*acetylase was nonfunctional. It is known that *O*-acetylation blocks the collective activity of all LTs and compounded by the dysfunction of Ape1 this could explain the pro-inflammatory potential of the Δ*ltgA*^*ltgA*Δ30^ mutant. Furthermore, the Δ*ltgA*^*ltgA*Δ30^ mutant bacteria form aggregates, potentially rendering them more susceptible to killing by phagocytes, and therefore cleared from the host more quickly than the other strains.

The initial innate host response to a *N. meningitidis* systemic infection is mediated by LPS sensing through TLR4, which would be the same for WT and Δ*ltgA* mutant. Therefore, the WT and Δ*ltgA* strains would have the same pro-inflammatory potential. However, clearing of the *N. meningitidis* infection is the result of effective activation of phagocytes such as neutrophils and it has been shown that efficient activation of neutrophils requires Nod1 sensing^60, 61^. Impaired release of monomeric muropeptides by the Δ*ltgA* mutant strain, resulting in reduced Nod1 activation, could explain why the Δ*ltgA* strain is cleared more slowly than the wild-type bacteria. Overall, these finding suggests that LtgA and possibly other LTs, plays a primary role in the life cycle of the cell, and regulate the function of other PG glycan strand specific enzymes that ensures the proper function of the cell wall machinery.

In summary, a mechanistic role for LtgA and other LTs emerges, whereby together with other PG modifying (LtgE, LtgD, Ape 1) or polymerization (PBP 1a) enzymes, they form the nucleus of the PG glycan strand remodeling machinery that is essential for the insertion of new material during growth. Ape1, a PG-de-*O*-acetylase, removes the *O*-acetyl group from the C6-hydroxyl position of the glycan strand (Fig. 8). Ape1’s function is regulated and stabilized by LtgA possibly because their functions are symbiotic. The removal of the *O*-acetyl group by Ape1 allows LtgA and LtgE to continue metabolism of the PG (Fig. 8). The endolytic enzyme (LtgE) likely cleaves internal sites on the glycan strands meanwhile, the exolytic enzyme (LtgA) cleaves the end of the PG glycan strands. The bifunctional PBP1a polymerizes the insertion of new material after remodeling or degradation (Fig. 8). Meanwhile, FtsN although not addressed experimentally in this paper likely connects LtgA to the cell division site and the PG polymerization machinery (FtsW-PBP2). An impaired LtgA disrupts the normal course of bacterial cell division or cell separation paving the way for the design of inhibitors or antibiotics that target LTs.

**Figure 8.**
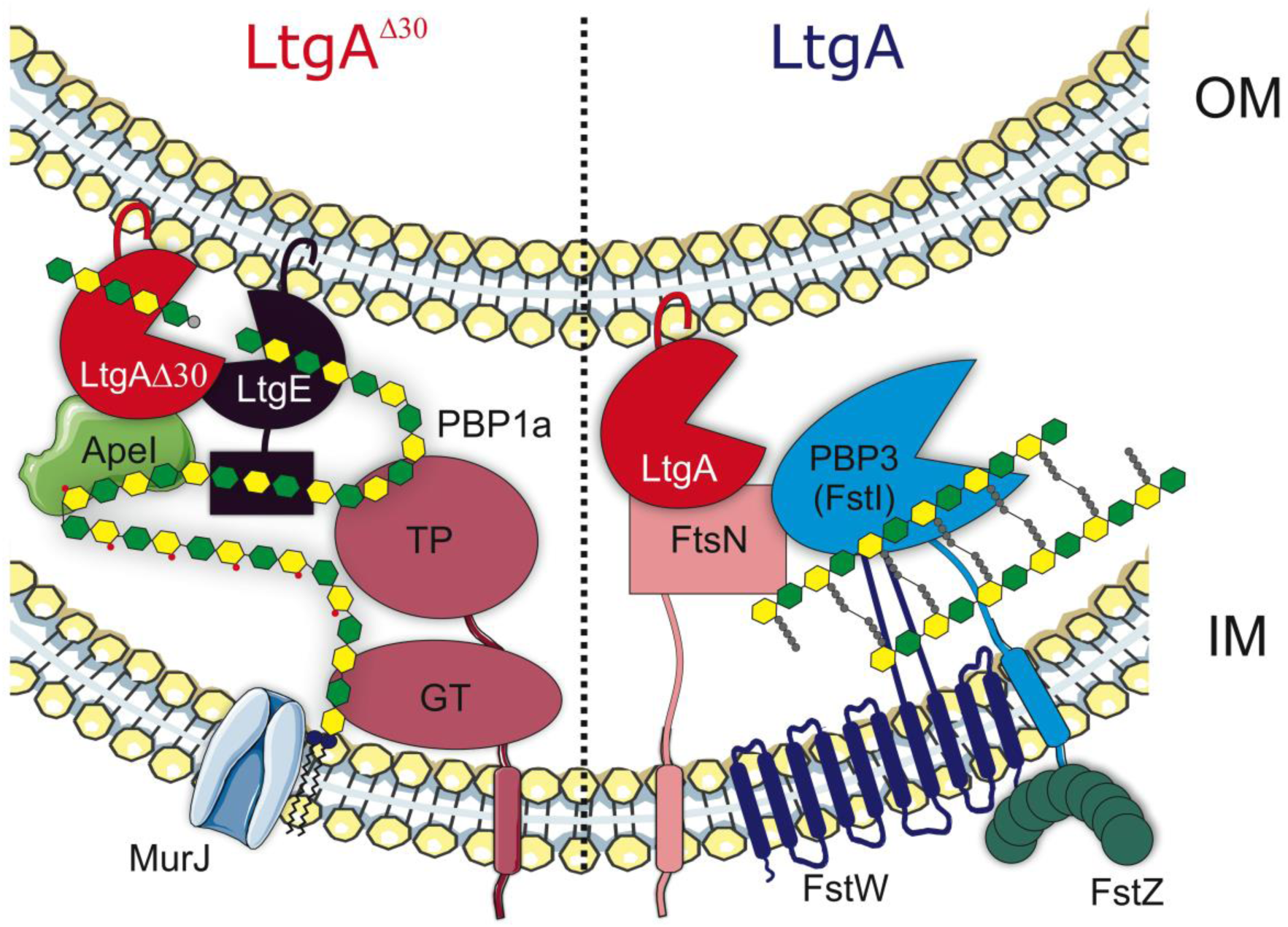
The proposed role of the essential PG centric protein hub directed by LtgA. The panel labelled LtgA^Δ30^ reveals the core of the glycan strand remodelling machinery, whereby Ape1 removes the *O*-acetyl group from the C6-hydroxyl position of the glycan strand. The removal of the *O*-acetyl group by Ape1 allows LtgA and LtgE to metabolize the PG. The endolytic enzyme (LtgE) cleaves nascent glycan strands and the exolytic enzyme (LtgA) cleaves old PG strands. The bifunctional PBP1a polymerizes the PG. The panel labeled LtgA decribed how FtsN connects LtgA to the PG synthesis at the cell division site.

## Conclusion

We devised a multidisciplinary approach using structural biology to show that it is possible to target a ‘hot spot’ on an LT in order to affect bacterial growth, cell division, and cell membrane integrity, which resulted in lethal consequences for the bacteria during host infection. Additionally, as we discovered with Ape1, LTs can regulate the function and activity of its binding partners. Our study further suggests that although the bacteria devised a safety net system of redundant LTs that were previously deemed non-essential to bacterial survival, we demonstrate that traditional ways of assessing their essentiality does not apply. However, inactivation of an LT primarily guided by structural knowledge of how it interacts with its substrates or protein partners helped to reveal its role in the bacterium. This study shows the ripple effects of disrupting LtgA PG binding capabilities and sets the stage for future development of a class of antibiotics that may act by a dual action *in vivo*. A small molecule binding to alpha helix 30 could interfere with growth and simultaneously promote bacterial clearance, mimicking the enhanced clearance of the Δ*ltgA*^*ltgA*Δ30^ mutant in a murine infection model.

## Methods

### Protein expression and purification

All constructs were created using standard molecular biological techniques. All constructs used for protein expression and purification in this study were GST fusions expressed from pGEX-4T1 (GE Life Sciences). The native proteins without signal peptides were expressed in BL21(DE3) Gold competent cells (Novagen). The gene encoding the LtgA deletion mutant lacking the alpha helix ^503^(ATAREIAGKIGMD)^513^ was chemically synthesized by ProteoGenix. The synthesized *ltgA* deletion gene was cloned into a GST-fusion pGEX-4T1 (GE Life Sciences) plasmid as described above. The expression of all constructs was induced with 0.6 mM IPTG at an optical density at 600nm (OD600) of 0.7-0.8 and harvested after 4 h of induction at 18°C. After glutathione-affinity chromatography and thrombin cleavage, proteins were purified to homogeneity by size-exclusion chromatography (Superdex-200, GE) in 50 mM HEPES (pH 7.4), 150 mM NaCl, and 1 mM BME. After gel filtration, the proteins were immediately used for crystallization or flash frozen in liquid nitrogen and stored at −80°C.

### X-ray crystallography

Crystallization screening was carried out by the sitting-drop vapor-diffusion method with a Mosquito^®^ (TTP Labtech) automated crystallization system. All crystals were grown at 18°C using the hanging-drop vapor-diffusion method. Crystals of 15-20 mg/ml LtgA were grown at 18°C and appeared within 2-3 days. LtgA was crystallized in a 1:1 (v/v) ratio against a well solution of 33% (w/v) PEG 6000 and 100 mM HEPES, pH 7.5. Crystals were rectangular in shape and grew to approximately 200-300 μm in length.

The data set was collected at the Soleil Synchrotron (Beamline Proxima-1) (Supplementary Table 3). Phasing by molecular replacement was performed using Phenix^62^. Building was performed using Coot^63^, and restrained refinement was carried out using a combination of Phenix and the ccp4 software suite^62,64^. MolProbity was used during building and refinement for iterative structure improvements^65^.

All structural figures were generated with PyMOL (PyMOL Molecular Graphics System, version 1.5, Schrödinger, LLC). The crystallographic parameters, data statistics, and refinement statistics are shown in Supplementary Table 3. Modeling of unknown LTs were accomplished using Phyre 2^66^. Movies of the LtgA enzymatic steps was generated in PyMOL and then assembled in photoshop and edited in iMovie.

### Protein-protein interaction studies by gel filtration

To explore the interactions of LtgA and its PG binding partners, proteins were mixed at equimolar concentrations of 10 µM, incubated on ice for 1 h, and subjected to gel-filtration chromatography on an SD200 10/300 column. Approximately 150-300 µl of each sample was applied to the column in 50 mM HEPES (pH 7.5) and 150 mM salt. Peak fractions were then subjected to analysis 5-15% SDS-PAGE.

### Analysis of Neisseria sp. peptidoglycan by reversed-phase HPLC and mass spectrometry

The peptidoglycan isolated from all four strains peptidoglycan (wild-type, *ΔltgA*, Δ*ltgA*^*ltgA*^ and Δ*ltgA*^*ltgA*Δ30^) was incubated for 16 h in the presence of 10 µg of mutanolysin in 12.5 mM sodium phosphate buffer (pH 5.8) at 37°C (total reaction volume 150 µl). The reaction was stopped by boiling the samples for 3 min, and the supernatant containing the soluble muropeptides was collected after centrifugation at 16,000 × *g* for 10 min. The supernatant was analyzed by reversed-phase HPLC using a Hypersil GOLD aQ column (5 μm particle size, 150 × 4.6 mm, flow rate 0.5 ml at 52°C, Thermo Fisher Scientific) with a mobile phase of H_2_O-0.05% trifluoroacetic acid and a 25% acetonitrile gradient over 130 min. Muropeptides of interest were collected and identified by mass spectrometry as previously described ^4,5^.

### Bacterial strains

Clone 12 is a derivative of strain 8013, a serogroup C *N. meningitidis* strain^67^, and MC58 is a serogroup B strain^68^. Bacteria were grown on GCB medium (Difco) containing Kellogg’s supplements^69^. The *E. coli* strain DH5^70^ was used for plasmid preparation and subcloning. Kanamycin, ampicillin and erythromycin were used in *E. coli* at final concentrations of 50, 100 and 300 µg/ml, respectively. In *N. meningitidis*, kanamycin, ampicillin and erythromycin were used at final concentrations of 100, 20 and 2 µg/ml, respectively.

### Plasmid construction

The *ltgA* gene (1851 nucleotides according to the genome sequence of the meningococcal strain MC58) was chemically synthesized with a deletion of 42 bp (14 codons) between positions 1506 (codon 502) and 1548 (codon 516) (starting at ATG) and was cloned into the vector pUC57 to generate the recombinant plasmid pUC57ltgA (ProteoGenix, Schiltigheim, France). The *ltgA* fragment was amplified using the primer pair NMF1/NMR1 from the plasmid pUC57ltgA and from the strain MC58. The two fragments were blunt-ended using the Klenow DNA polymerase fragment (BioLabs) and subcloned into the *BamH*I site of the recombinant plasmid pTE-KM^71^. This plasmid contains the *pilE* gene of clone 12 with the Km cassette, encoding resistance to kanamycin, located immediately downstream of the *pilE* without modification of *pilE* expression. Moreover, a unique *BamH*I site located between the Km cassette and the downstream region at the 3’ end of the *pilE* gene^67,71^ was used to subclone the two blunt-ended fragments from the plasmid pUC57ltgA and from the strain MC58 to yield the recombinant plasmids pD-Δ*ltgA*^*ltgA*^ and pD-Δ*ltgA*^*ltgA*Δ30^, respectively.

An internal deletion in the *ltgA* gene was also constructed by removing the segment between the restriction sites *BsmI* (position 21) and *BalI* (position 1724) on pUC57ltgA, and this region was replaced with the *ermAM* cassette, encoding erythromycin resistance; the construct was checked using the primer pair ERAM1/ERAM3 (5’-gcaaacttaagagtgtgttgatag-3’ and 5’-aagcttgccgtctgaatgggacctctttagcttcttgg-3’, respectively)^71^. The corresponding recombinant plasmid pDG15-09 was linearized at the *EcoR*I site of the pUC57 vector and used to transform the clone 12 strain of *N. meningitidis*. Transformants were selected on standard GCB medium in the presence of 2 µg/ml erythromycin. Integration by homologous recombination into the *ltgA* gene on the meningococcal chromosome was further confirmed by PCR analysis using the oligonucleotides ERAM1/ERMA3 and NMF1/NMR1. One transformant was selected for further analysis and named pD-ΔltgA.

The two recombinant plasmids pD-Δ*ltgA*^*ltgA*^ and Δ*ltgA*^*ltgA*Δ30^ were linearized using the ScaI restriction enzyme and used to transform the strain pD-*ΔltgA*. Transformants were selected on standard GCB medium in the presence of 2 µg/ml erythromycin and 100 µg/ml kanamycin. Integration by homologous recombination into the *ltgA* gene on the meningococcal chromosome was further monitored by PCR analysis using the oligonucleotides pilE1, NMF1, NMR1 and NMF1/NMR1. One transformant from each transformation was selected for further analysis and named Δ*ltgA*^*ltgA*^ or Δ*ltgA*^*ltgA*Δ30^. The strain Δ*ltgA*^*ltgA*^ has the *ltgA* gene deleted from its locus but harbors the *ltgA* gene downstream of the *pilE* site. The Δ*ltgA*^*ltgA*Δ30^ strain also has the *ltgA* gene deleted from its locus and contains a downstream *pilE* gene but harbors the *ltgA* gene with the region encoding the amino acid residues 501-516 deleted.

### Coimmunoprecipitation studies

Coimmunoprecipitation (IP) studies were carried out with lysates prepared from the *N. meningitidis* MC58 strain. Harvested cells were washed in ice-cold 1× PBS. Cells were lysed in a solution containing 20 mM Tris (pH 8), 137 mM sodium chloride, 1% Nonidet P-40 (NP-40) detergent, 1 mM EDTA and protease inhibitor cocktail (1 tablet/20 ml) to obtain approximately 1×10^7^ cells per ml. Cells were incubated for 2 h at 4°C and then microcentrifuged for 15 min at 12,000 rpm. The supernatant was carefully removed into a fresh tube and kept on ice. The lysate was then precleared to prevent nonspecific binding to the beads. Briefly, 100 μl of freshly prepared protein A sepharose beads was incubated with 1 ml of lysate for 2 h at 4°C with gentle agitation. This slurry was then centrifuged for 10 min at 14,000 ×*g*. Pelleted beads were discarded, and the supernatant was recovered. In a microcentrifuge tube, 50 μg of the lysate was incubated overnight with 5 μl of a polyclonal antiserum to LtgA at 4°C under gentle agitation. Seventy microliters of beads was added to each sample and incubated at 4°C for 4 h. Samples were then centrifuged at 14,000 rpm, and the supernatant was discarded. The beads were then gently washed by centrifugation at 4°C in lysis buffer three times to eliminate nonspecific binding. Fifty microliters of the sample was heated at 90°C for 5 min in 2× loading buffer, loaded onto an SDS gel and analyzed by western blotting.

### Immunoprecipitation and LC-MS/MS analysis

For immunoprecipitation of LtgA from Δ*ltgA*^*ltgA*^, Δ*ltgA*^*ltgA*Δ30^, and *ΔltgA*, beads were washed 4 times with 1 ml of ice-cold PBS and 4 times with 1 ml of ice-cold digestion buffer (20 mM Tris (pH 8.0), 2 mM CaCl_2_). The washed beads were resuspended in 200 µl of digestion buffer and incubated for 5 h with 1 µg of trypsin (Promega) at 37°C. The beads were removed, an additional 1 µg of trypsin was added, and the proteins were further digested overnight at 37°C. Peptides were purified by OMIX C18 tips (Agilent), dried and redissolved in 30 µl of 0.1% formic acid in water/acetonitrile (98:2, v/v), of which 1 µl was injected for LC-MS/MS analysis on an EASY-nLC 1000 system (Proxeon, Thermo Fisher Scientific) connected in line to a Q Exactive Plus mass spectrometer with a Nanospray Flex ion source (Thermo Fisher Scientific). Peptides were loaded in solvent A (0.1% formic acid in water) on a reversed-phase column (made in-house, 75 µm i.d. × 500 mm, 1.9 µm beads, C18 Reprosil-Pur Dr. Maisch) and eluted by increasing solvent B (0.1% formic acid in acetonitrile) in linear gradients from 5% to 27% in 100 min, then from 27% to 45% in 40 min, and finally from 45% to 60% in 10 min, all at a constant flow rate of 250 nl/min. The mass spectrometer was operated in data-dependent mode, automatically switching between MS and MS/MS acquisition for the five most abundant ion peaks per MS spectrum. Full-scan MS spectra (300-1700 m/z) were acquired at a resolution of 70,000 after accumulation to a target value of 3,000,000 with a maximum fill time of 20 ms. The five most intense ions above a threshold value of 170,000 were isolated (window of 1.6 Th) for fragmentation at a normalized collision energy of 27% after filling the trap at a target value of 1,000,000 for a maximum of 60 ms with an underfill ratio of 1%. The S-lens RF level was set at 60, and we excluded precursor ions with single, unassigned and charged states greater than six from fragmentation selection.

### Fluorescent labeling and fluorescent microscopy

Bacterial cultures were centrifuged 5 minutes at 5000 rpm and re-suspended in PBS containing 1 µg/mL DAPI and 5µg/ml FM4-64 FX (*N*-(3-Triethylammoniumpropyl)-4-(6-(4 (Diethylamino) Phenyl) Hexatrienyl) Pyridinium Dibromide) probe. The cells were incubated for 10 minutes at room temperature protected from light, centrifuged and the pellets resuspended in 4% pFA for fixation during 5 minutes. After fixation, the cells were washed with PBS, and a 10 µL drop of the bacterial suspension was applied onto poly-Lysine pre-coated cover glasses (# 1.5). Next, samples were mounted using Prolong Diamond and imaged using Leica SP5 confocal microscope, with a 63X (1.4 NA) oil-immersion objective using 405 nm and 514 nm laser lines. Fluorescence was recorded sequentially using hybrid (HyD) detectors and images processed using Fiji^72^.

### Scanning electron microscopy

*Neisseria meningitidis* were prefixed in 2.5% Glutaraldehyde diluted in PHEM (Pipes, Hepes, EGTA and MgSO_4_) buffer at pH 7. The cells were prefixed for 1 hr at room temperature, followed by 2 washes in PHEM buffer. The samples were applied onto the cover glass (1.5 mm) pre-coated with poly-Lysine. This was followed by a light speed centrifugation to ensure that the cells adhere correctly to the cover slip.

The bacteria were post-fixed using 2% osmium tetroxide in PHEM buffer for 30 to 60 minutes followed by washing with water three times. The specimen was dehydrated using increasing ethanol concentrations of 25% to 100% in increments of 25%. The bacteria were critically point dried using carbon dioxide, coated with gold and examined with the JEOL JSM 6700F scanning electron microscope.

### Protein identification and quantification

Data analysis was performed with MaxQuant (version 1.6.3.4)^73^ using the Andromeda search engine with the default search settings, including a false discovery rate of 1% at both the peptide and protein levels. Spectra were searched against a database of 2204 proteins from *N. meningitidis* strain 8013 (taxid 604162, downloaded from http://www.ncbi.nlm.nih.gov/). The mass tolerance for precursor and fragment ions was set to 4.5 and 20 ppm, respectively, during the main search. Enzyme specificity was set as C-terminal to arginine and lysine, also allowing cleavage at proline bonds, with a maximum of two missed cleavages. Variable modifications were set to oxidation of methionine residues. Only proteins with at least one unique or razor peptide were retained, leading to the identification of 879 *Neisseria* proteins. Proteins were quantified by the MaxLFQ algorithm integrated in MaxQuant software^74^. A minimum ratio count of two unique or razor peptides was required for quantification. Further data analysis was performed with Perseus software (version 1.5.2.4) after loading the protein group file from MaxQuant LFQ intensity values were log2 transformed and replicate samples of the three strains were grouped. Proteins with less than three valid values in at least one group were removed, and missing values were imputed from a normal distribution around the detection limit. Then, t-tests were performed (FDR=0.05 and S0=1) for pairwise comparisons of Δ*ltgA*^*ltgA*^ with *ΔltgA* and Δ*ltgA*^*ltgA*Δ30^ with ΔltgA. The results of these t-tests are listed in Supplementary table 2.

### *O*-acetyl peptidoglycan esterase assay

The acetyl esterase activity assays were executed as previously described with minor modifications^42,75^. Briefly, the reaction utilized 2 mM 4-nitrophenyl acetate as the substrate. The reaction was carried out at 37 °C in 50 mM sodium phosphate buffer, pH 6.5 in the presence of LtgA, using equimolar amounts of LtgA and Ape1 or Ape1. The final volume of the reaction was 300 μl. The reaction was initiated with the addition of the substrate 4-nitrophenyl acetate dissolved in 5% v/v ethanol. The release of 4-nitrophenyl was monitored over the time course of an hour in 96 well microtiter plate at an absorbance of 405 nm.

### Analysis of LtgA activity

To assess the activity of LtgA, PG (200 µg) was incubated in the presence of LtgA, or equimolar amounts of LtgA and Ape1, in 12.5 mM sodium phosphate buffer pH 5.6. *Neisseria* PG was purified as previously described^76^. The reaction mix was initiated by the addition of enzymes and incubated at 37 °C for 5 min. Control reactions lacking PG or enzyme/inhibitor were also included. The final reaction volume was 200 µL. Reactions were performed in triplicates. The reaction was stopped by incubating the samples in a heat block at 100 °C for 5 min. The soluble 1,6-anhydro-muropeptides was collected using centrifugation at 16,000 *g* for 10 min at room temperature. The supernatant was collected and analyzed by reversed-phase HPLC using a Shimadzu LC-20 system with a Hypersil GOLD aQ column (5 μm particle size, 250 × 4.6 mm, flow rate 0.5 mL/mL at 52 °C; Thermo Fisher Scientific (Waltham, MA, USA). The mobile phase gradient was 50 mM sodium phosphate pH 4.3 to 75 mM sodium phosphate pH 4.9 with 15% Methanol over 135 min.

### Infection model

A previously published model for meningococcal infection in transgenic mice expressing human transferrin was used ^77^. Four strains were tested: clone 12 (wild-type), Δ*ltgA*^*ltgA*^, Δ*ltgA*^*ltgA*Δ30^ and *ΔltgA*. Five mice per group were infected by intraperitoneal injection with 500 µl of bacterial suspension of each strain at 1×10^7^ CFU/ml. Blood samples were obtained by retro-orbital bleeding after 2, 6 and 24 h, and bacterial counts were determined by plating serial dilutions on GCB medium.

### Phylogenetic tree construction

Protein sequences were aligned using MUSCLE alignment algorithm using UPGMA clustering method in MEGAX^78^. Using aligned sequences, a maximum likelihood tree was constructed using a Neighbor joining construction method and a JTT protein substitution model in CLC Genomics Workbench 8.01. Robustness was estimated using 500 bootstrap replicates (values not shown in figures).

#### Ethics Statement

Animal work in this study was carried out at the Institut Pasteur in strict accordance with the European Union Directive 2010/63/EU (and its revision 86/609/EEC) on the protection of animals used for scientific purposes. The laboratory at the Institut Pasteur has the administrative authorization for animal experimentation (Permit Number 75-1554) and the protocol was approved by the Institut Pasteur Review Board that is part of the Regional Committee of Ethics of Animal Experiments of Paris Region (Permit Number: 99-174). All the invasive procedures were performed under anesthesia and all possible efforts were made to minimize animal suffering.

### Cytokine assay

Blood samples from infected mice were collected and stored at −80°C. Cytokines (IL-6 and KC) were quantified by an enzyme-linked immunosorbent assay (Quantikine; R&D Systems Europe, Abingdon, Oxon, United Kingdom).

### Growth curves and LtgA stability assay

Bacteria were grown overnight in GC broth with Kellogg’s supplements at 37°C and 5% CO_2_. Fresh medium was inoculated at an OD600 of 0.05, and growth was measured spectrophotometrically at 1-h intervals over a period of 24 h at 37°C and 5% CO_2_. When indicated, 2 µg/ml chloramphenicol was added when the OD_600_ reached 0.6, and incubation was continued at 37°C and 5% CO_2_. At different incubation time points, aliquots (3 ml) from each culture were sampled, and the bacteria were collected by centrifugation, lysed by boiling in SDS-containing sample buffer, and analyzed for the presence of LtgA by western blotting using anti-LtgA antibodies. The expression of the outer membrane factor H binding protein (Fhbp) was used as an internal control.

### PG binding assay

The binding of the different LtgA proteins to PG was carried out by incubating 100 µg of PG and 10 µg of enzymes suspended in 150 µl of Tris buffer pH 7.5 (10 mM Tris, 10 mM MgCl_2_ and 50 mM NaCl). After 30 min of rocking at room temperature, 50 µl of the sample was set aside for analysis before centrifugation for 10 min at 20,000 x*g*. The supernatant was discarded, and the insoluble fraction was washed three times. The remaining pellet was boiled for 10 min. Five microliters of the input or unbound and bound fractions was loaded on an SDS-PAGE gel and analyzed by western blotting.

### Accession numbers

Coordinates and structural data have been submitted to the Protein Data Bank under the accession code 6H5F. The mass spectrometry proteomics data have been deposited to the ProteomeXchange Consortium via the PRIDE partner repository with the dataset identifier PXD013393 (reviewer login; password: mkJ4Yxxn).

## Acknowledgments

We would like to acknowledge the beamline staff (PROXIMA-1 at SOLEIL and X06DA at SLS) for their assistance. We are extremely grateful to Frederick Saul, Patrick Weber and Marco Bellinzoni for their constant helpful guidance, advice and assistance. We thank Dr. Antoine Forget for the advice on and help with figure presentations. A.H.W. was supported by an EMBO long-term fellowship (ALTF 732-2010) and an Institut Carnot-Pasteur Maladies Infectious fellowship. This work was supported by an ERC starting grant (PGN from SHAPE to VIR 202283) and a Fondation pour la recherche médicale (FRM) grant Programme d’Urgence (DBF20160635726) 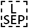 to I.G.B. This study received funding from the French Government′s Investissement d′Avenir program, 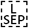 Laboratoire d′Excellence “Integrative Biology of Emerging Infectious Diseases” (grant n°ANR- 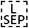 10-LABX-62-IBEID). F.I. was supported by a Pasteur Roux fellowship P.A.D.B. is supported by the European Union’s Horizon 2020 research and innovation program under the Marie Sklodowska-Curie grant agreement N° 665807 and by the Institut Carnot Pasteur Microbes & Santé. IS was supported by the Institut Carnot Pasteur Microbes & Santé given to the Pasteur-Paris University PhD program and the “Fin de these de science” number FDT201805005258 granted by “Fondation pour la recherche médicale (FRM).

## Contributions

A.H.W. and I.G.B. designed the research. A.H.W. purified and biochemically characterized the protein. R.W. and A.H.W. conducted the HPLC-based enzyme assays. A.H.W. and A.H. conducted all the structural experiments. A.H. collected the X-ray data. A.H.W. solved and refined all the structures. M-K.T., A.E.D, I.S. conducted biological validation of the structural studies. F.I., C.M.J.C-R, P.B. conducted mass spectrometric analysis. A.H.W., R.W., I.G.B., analyzed the data. W.A.H.W. wrote the manuscript. I.S., R.W., I.G.B., A.H., M-K.T contributed to the editing of the article.

## Supplemental Materials

**Supplementary Figure 1:**
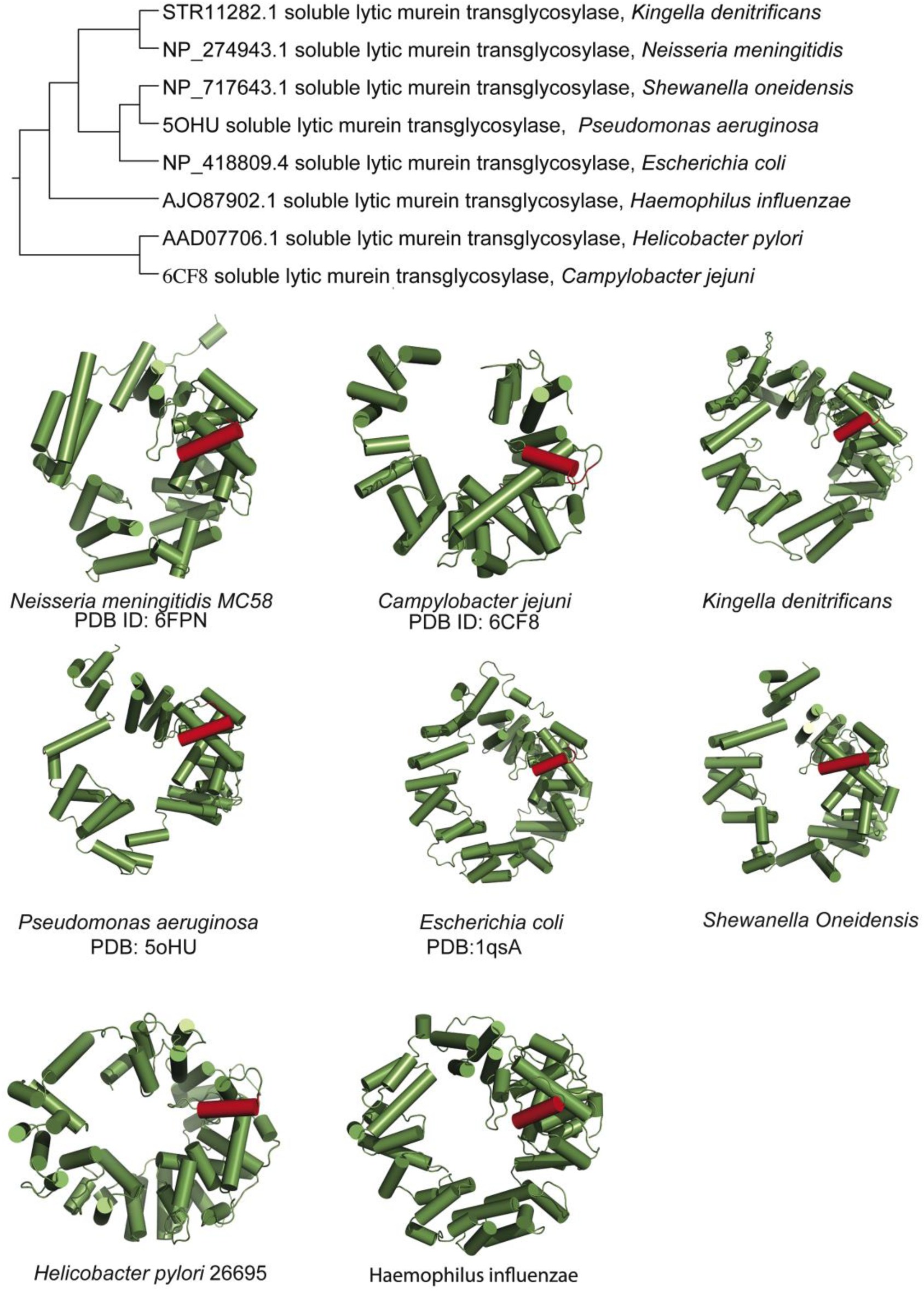
Conservation of alpha helix 30 amongst diverse lytic transglycosylases. Phylogenetic tree of lytic transglycosylases from various organisms complemented with various structures or predicted structures of lytic transglycosylases highlighing the conserved alpha helix 30 (PDB: protein data bank).

**Supplementary Figure 2:**
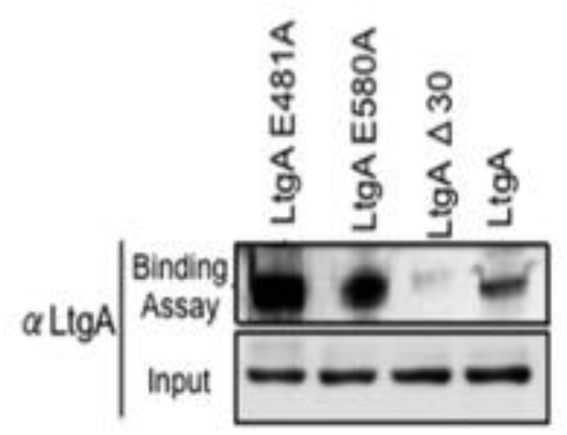
Binding of LtgA to Peptidoglycan. Heterologously expressed purified proteins of LtgA E481, LtgA E508A, and LtgA^Δ30^ were tested for their ability to bind *Neiserria* PG. Equal concentrations of purified protein (5 μg) were mixed with *Neisseria* PG and subjected to high-speed centrifugation. The western blot reflects proteins bound to insoluble PG. Comparatively, LtgA^Δ30^ appears to be defective in PG binding.

**Supplementary Figure 3.**
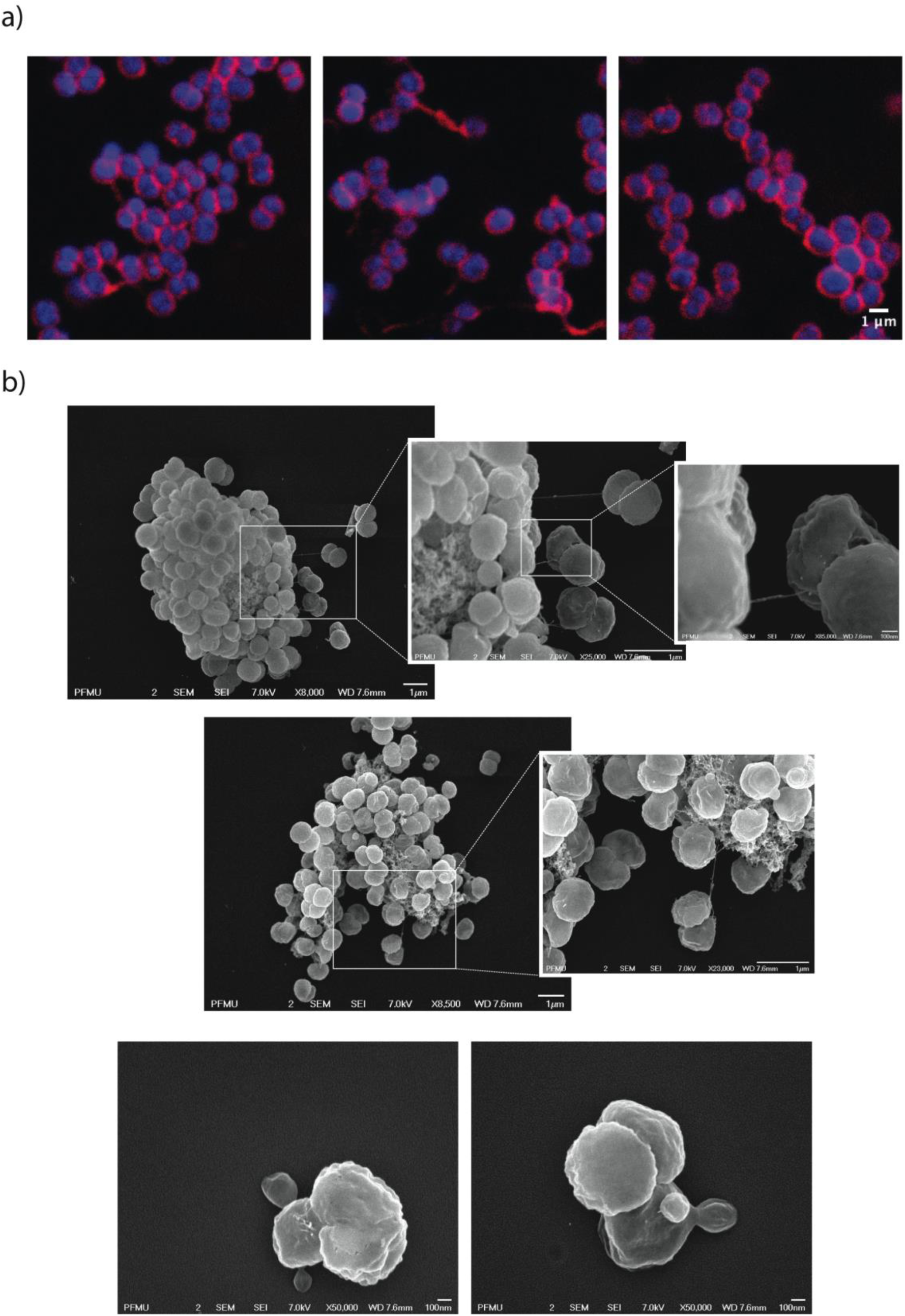
Morphological abnormalities of LtgA alpha helix 30 mutant. Fluorescent microscopy of Δ*ltgA*^*ltgA*Δ30^ strain highlighting aggregated cells that are defective in division and separation. b) Scanning electron microscopy of Δ*ltgA*^*ltgA*Δ30^ strain detailing additional morphological abnormalities such as, aggregation of extracellular material that resembles type IV pilin protein structures that stretches between diplococcic bacteria.

**Supplementary Figure 4.**
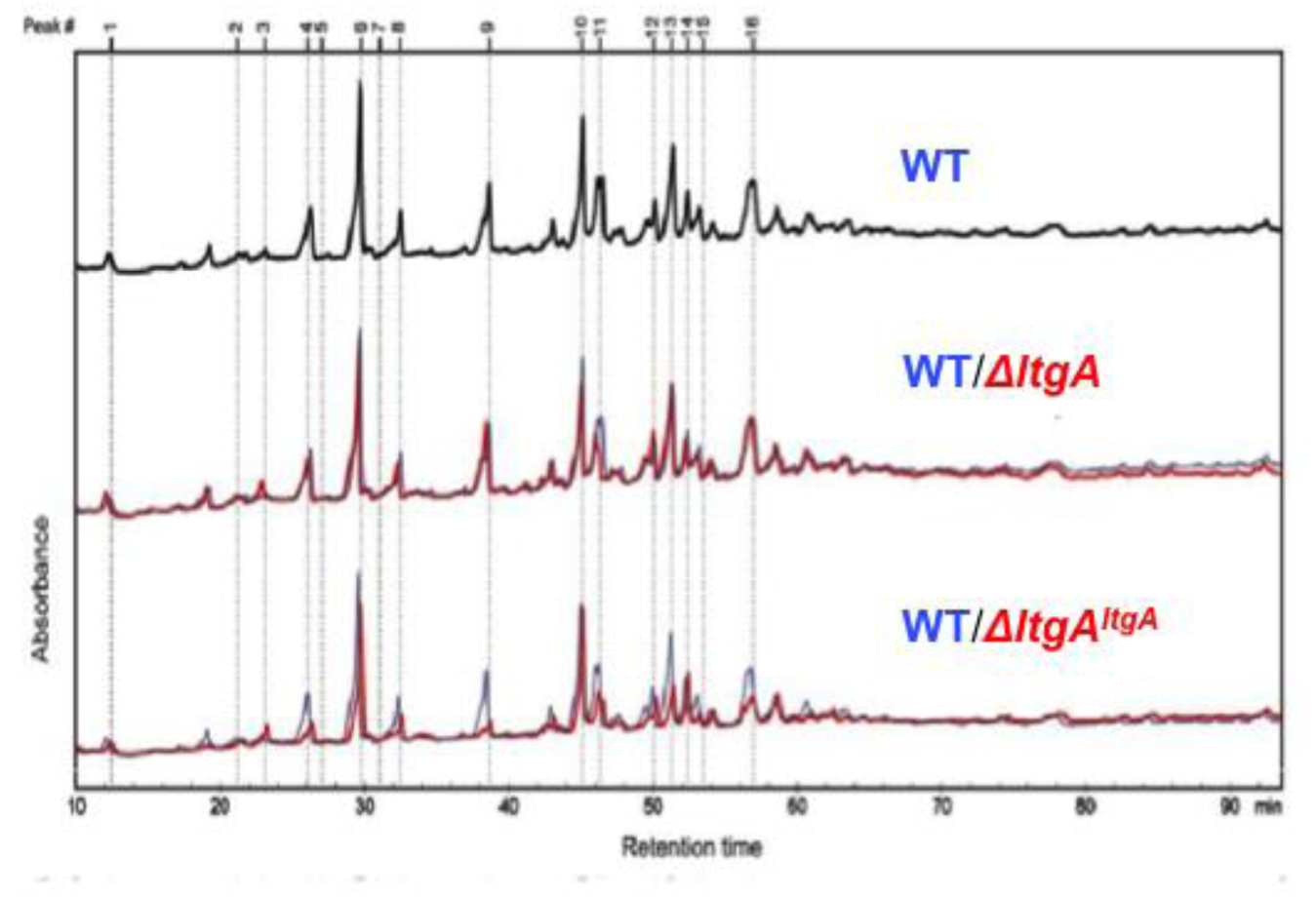
Muropeptide composition of PG isolated from wild-type, *ΔltgA, ΔltgA*^*ltgA*^. The purified PG was digested by muramidase mutanolysin, and the resulting muropeptides were reduced and then analyzed by LC/MS. The results were reproducible over 4 biological replicates. The wild-type chromatogram (blue) is overlaid on mutant chromatograms (red). Peak numbers correspond to Supplementary Table 1.

**Supplementary Figure 5.**
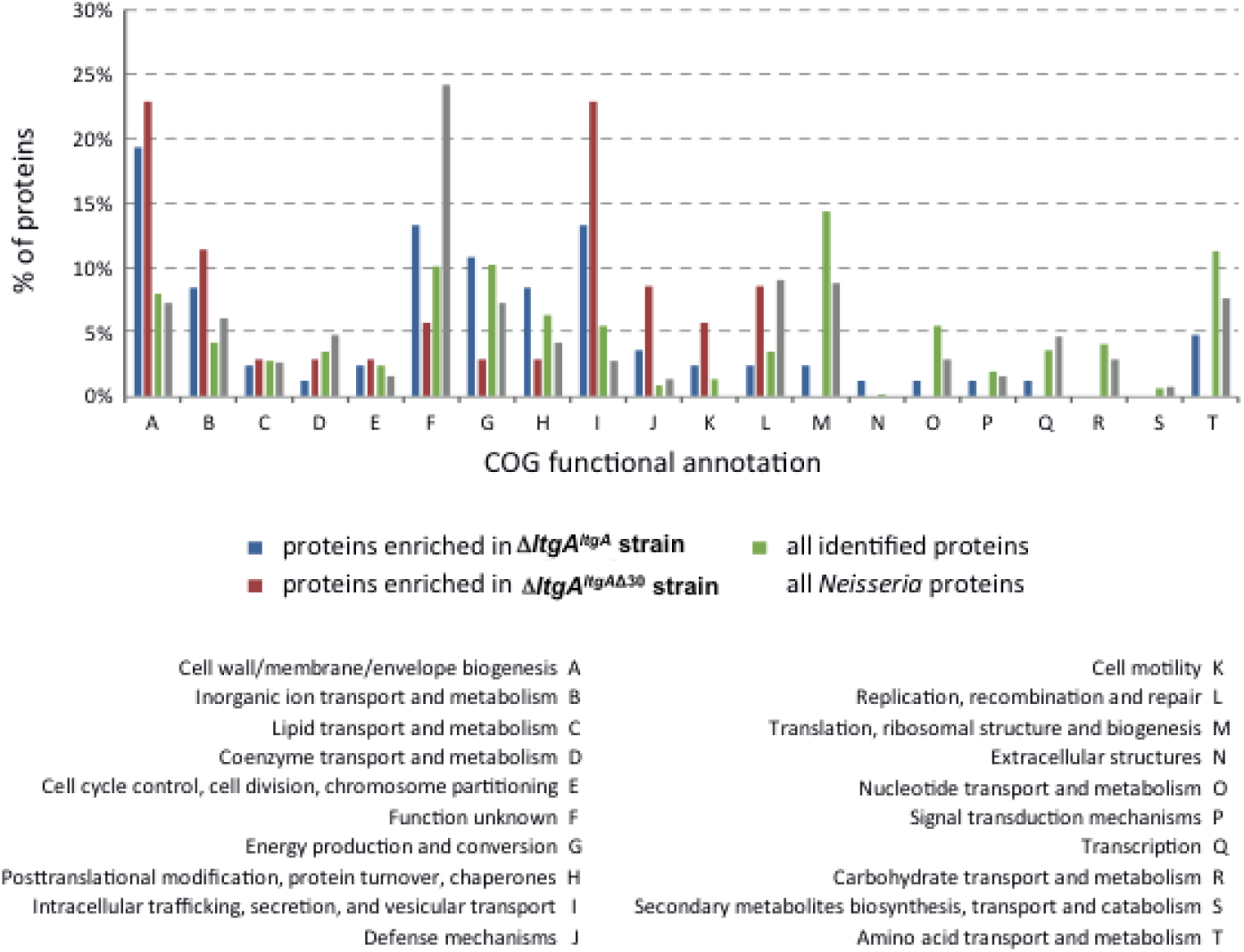
Clusters of orthologous groups of proteins (COG) analysis showed that the PPI network for LtgA^Δ30^ mainly occurred in two functional categories: cell wall/membrane/envelope biogenesis and intracellular trafficking, secretion, and vesicular transport.

**Supplementary Figure 6.**
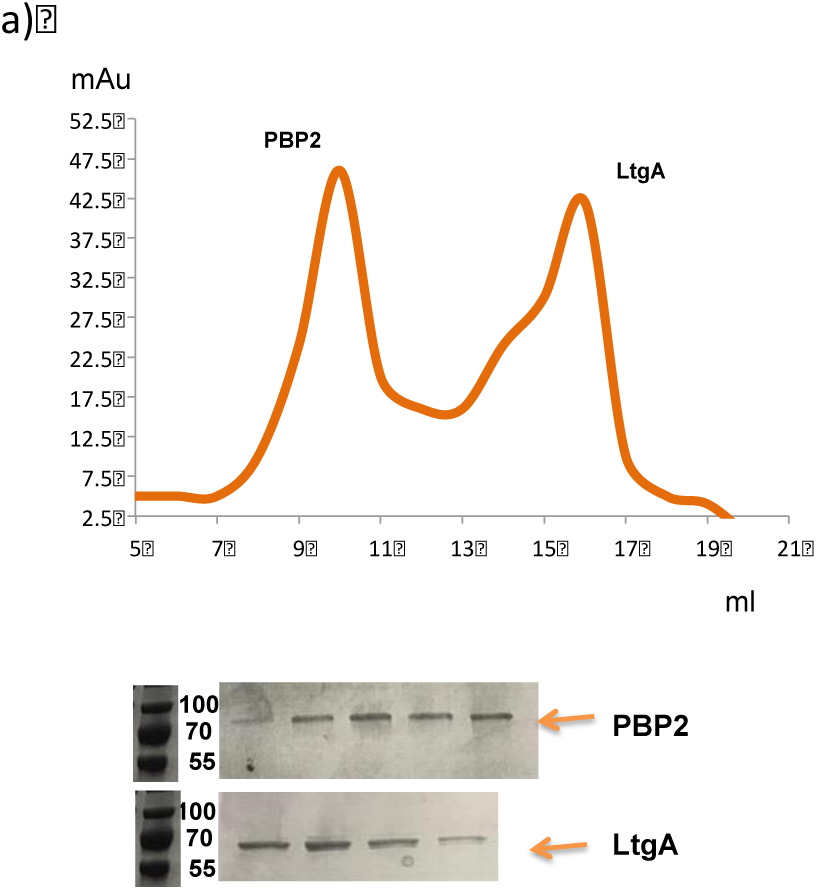
Gel filtration analysis of PBP2 and LtgA. Oligomeric PBP2 and LtgA run as separate species over gel filtration. Fractions were analyzed by using SDS-PAGE.

**Supplementary Figure 7.**
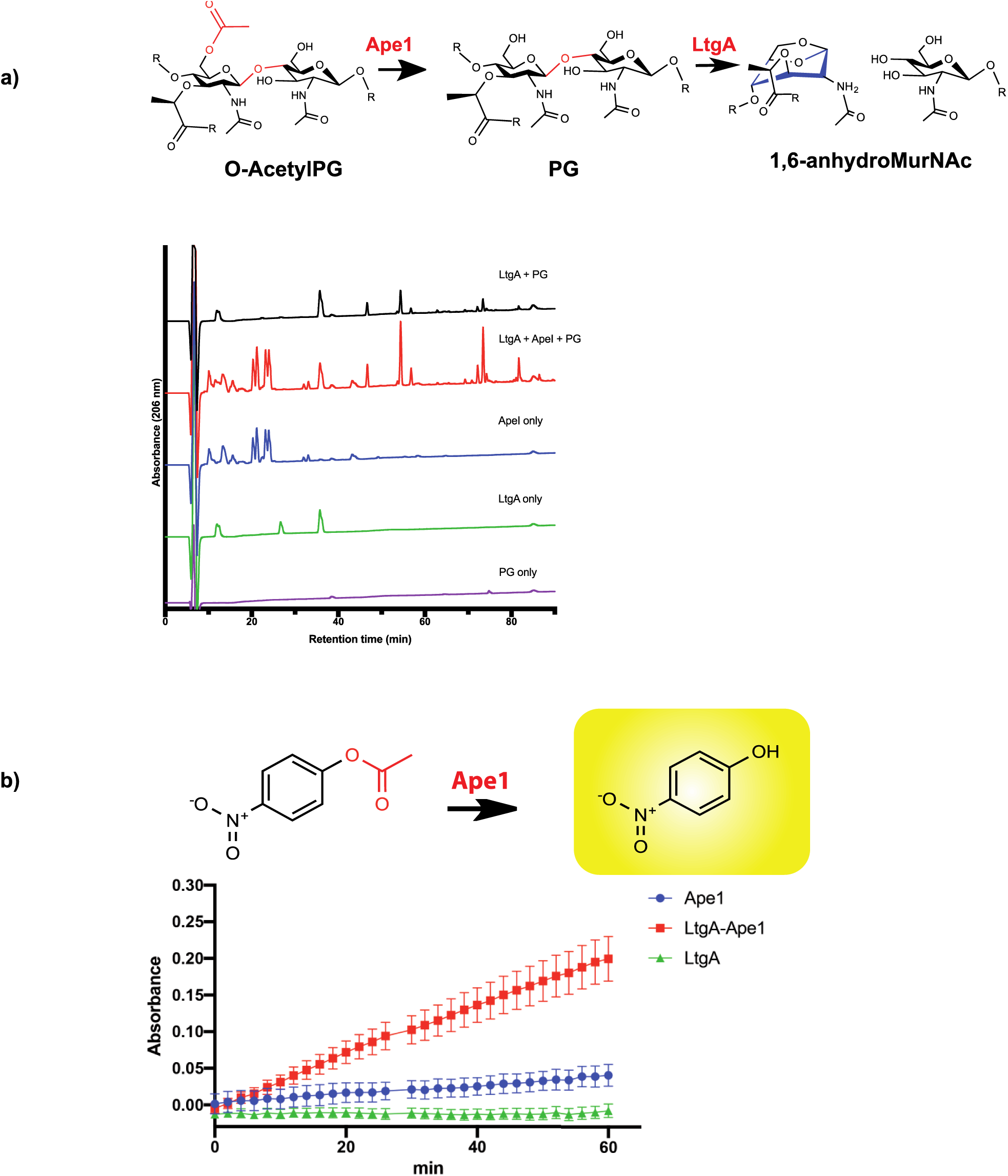
LtgA stimulates and stabilizes the enzymatic activity of Ape1. a) Stimulation of LtgA processivity toward acetylated PG by Ape1. b) LtgA stimulates and stabilizes Ape1 activity. Error bars shows the standard deviation of triplicates.

**Supplementary Figure 8.**
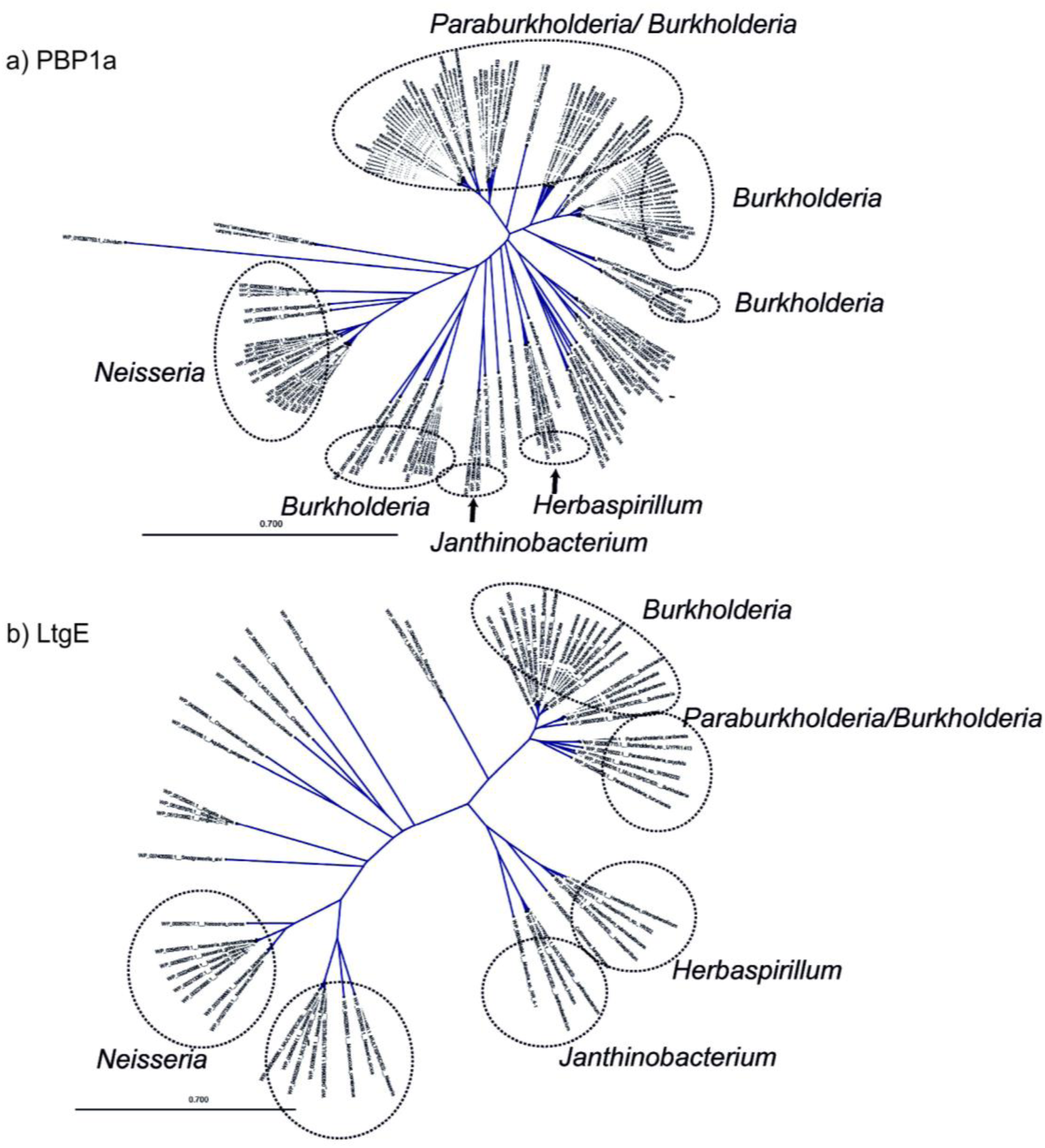
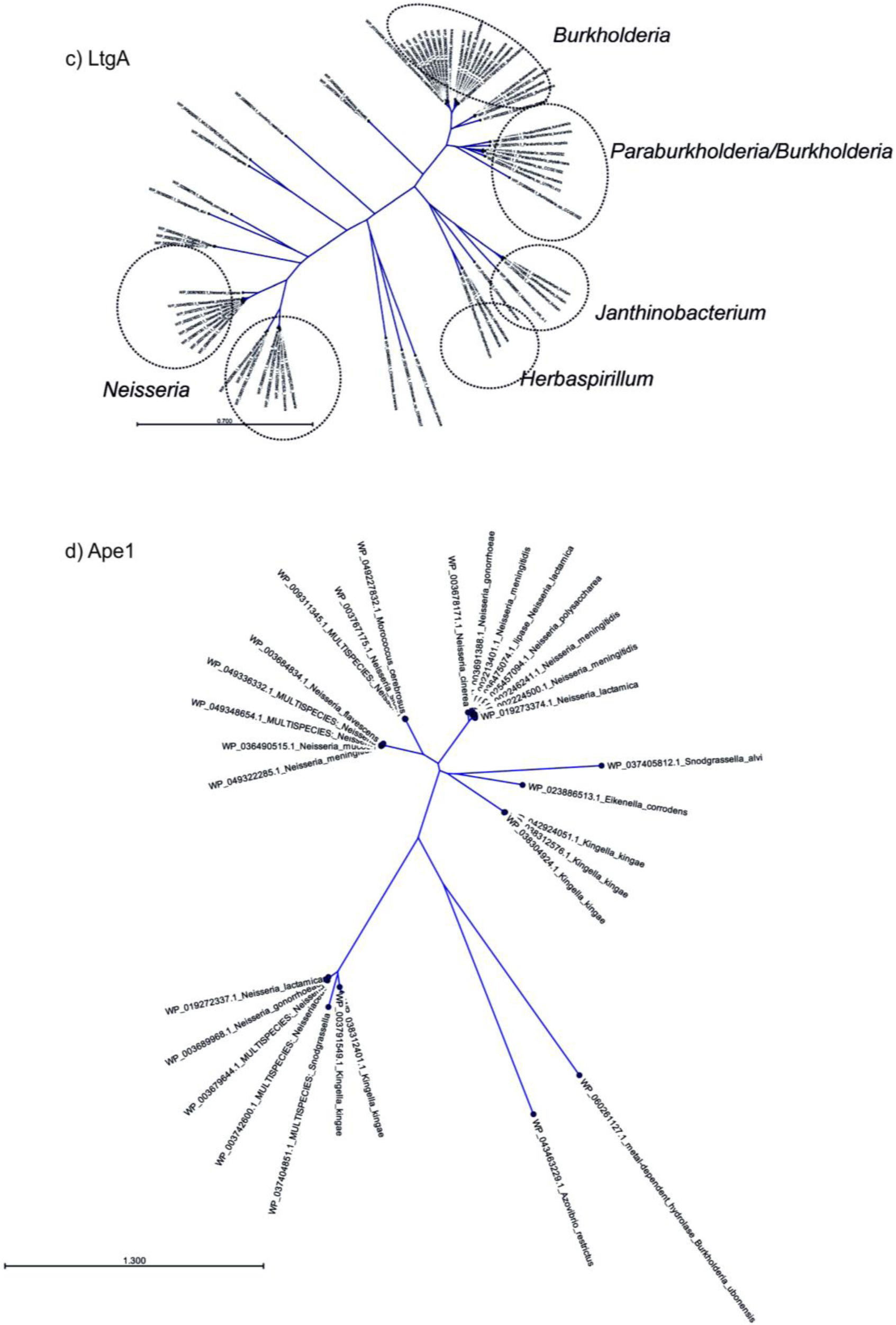
a-d) Phylogenetic tree showing PBP1a, LtgA, LtgE and Ape1 co-conservation.

**Supplementary Table 1.**
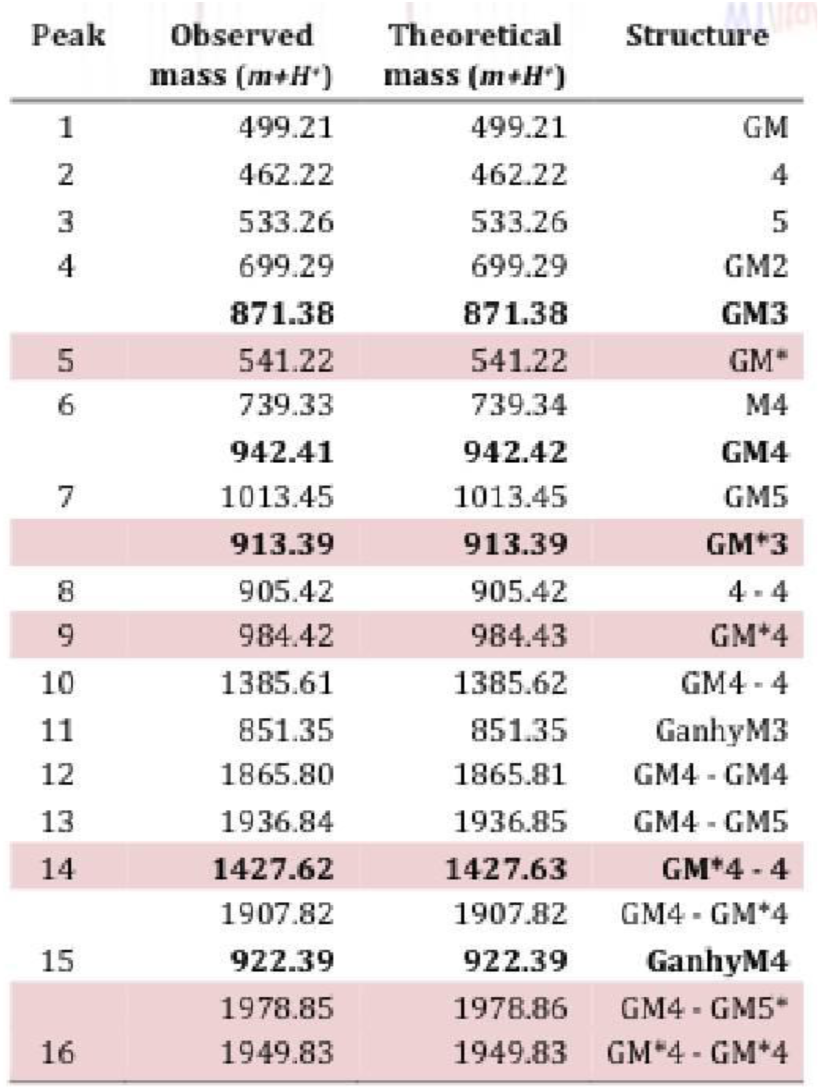
Muropeptides identified by mass spectrometry. * indicates *O*-acetylated MurNAc. Acetylated muropeptides are highlighted in pink. Where multiple muropeptides coeluted as a single peak, bold text indicates the most abundant mass detected.

**Supplementary Table 2. List of *Neisseria* proteins identified and quantified by LC-MS/MS after co-immunoprecipitation of LtgA.** Proteins that were identified as LtgA interaction partners are indicated in green (73 interaction partners were identified in the *ΔltgA*^*ltgA*^ strain and 34 interaction partners in the *ΔltgA*^*ltgA*Δ30^ strain). The values in column C indicate the fold change (in log2) between each pair of samples. The values in column D indicate the statistical significance (-log p value). For each protein, both values are also plotted in the volcano plots in Fig. 5 (fold change values on the x-axis and -log p values on the y-axis). See enclosed excel sheet.

**Supplementary Table 3.** Crystallography data collection and refinement statistics of LtgA.

|  |  |
| --- | --- |
| **Data Collection** | **LtgA** |
| **Ligand added** |  |
| Data collection |  |
| Wavelength (Å) | 0.9795 |
| Resolution range (Å) | 49.84-1.98 (2.067-1.995) |
| Space group | *P*2<sub>1</sub>2<sub>1</sub>2<sub>1</sub> |
| Unit-cell parameters |  |
| *a* (Å) | 66.82 |
| *b* (Å) | 72.25 |
| *c* (Å) | 122.31 |
| α(°),β(°),γ(°) | 90 |
| Total reflections | 299848 |
| Unique reflections | 40499 |
| Multiplicity | 7.4 (7.4) |
| Completeness (%) | 98.63 (94.17) |
| Mean *I*/σ(*I*) | 16.40 (1.47) |
| Wilson *B* factor (Å<sup>2</sup>) | 43.22 |
| *R*<sub>merge</sub><sup>#</sup> | 0.055 (0.063) |
| Refinement |  |
| *R* factor<sup>+</sup> | 0.1929(0.3192) |
| *R*<sub>free</sub> | 0.2464(0.3645) |
| No. of atoms | 4661 |
| No. of waters | 271 |
| No. of protein residues | 563 |
| R.m.s.d., bonds (Å) | 0.007 |
| R.m.s.d., angles (°) | 0.96 |
| Ramachandran favored (%) | 98 |
| Ramachandran outliers (%) | 0 |
| *B* factors (Å<sup>2</sup>) |  |
| Average | 51.00 |
| Macromolecules | 50.80 |
| Ligand | ----- |
| Solvent | 54.60 |
| All-atom clash score | 2.09 |

<sup>†</sup> $R_{\text{merge}} = \frac{\sum_{hkl} \sum_i |I_i(hkl) - \langle I \rangle_{(hkl)}|}{\sum_{hkl} \sum_i I_i(hkl)}$
<sup>‡</sup> $R_{\text{factor}} = \frac{\sum_{hkl} ||F_{\text{obs}}| - |F_{\text{calc}}||}{\sum_{hkl} |F_{\text{obs}}|}$
\**R*<sub>free</sub> was computed identically except that
all reflections belonged to a test set consisting of a 10% random selection of the data.

